# Dynamics of interdomain rotation facilitates FtsZ filament assembly

**DOI:** 10.1101/2022.10.13.512043

**Authors:** Joyeeta Chakraborty, Sakshi Poddar, Soumyajit Dutta, Vaishnavi Bahulekar, Shrikant Harne, Ramanujam Srinivasan, Pananghat Gayathri

## Abstract

FtsZ, the tubulin homolog essential for bacterial cell division, assembles as Z-ring at the division site, and directs peptidoglycan synthesis by treadmilling. To obtain insights into fundamental features of FtsZ assembly dynamics independent of peptidoglycan synthesis, we characterized the FtsZ from the cell wall-less bacteria, *Spiroplasma melliferum* (SmFtsZ). SmFtsZ was found to be a slower GTPase and has higher critical concentration (CC) for polymerization compared to *Escherichia coli* FtsZ (EcFtsZ). In FtsZs, a conformational switch from R (close)- to T (open)- state favors polymerization. In FtsZs, a conformational switch from R (close)- to T (open)- state favors polymerization. We identified a residue, Phe224, located at the cleft between N-terminal domain (NTD) and C-terminal domain (CTD) of SmFtsZ, which is crucial for R- to T-state transition. The mutation F224M in SmFtsZ cleft resulted in higher GTPase activity and lower CC, whereas the corresponding M225F in EcFtsZ resulted in cell division defects in *E. coli*. Our results demonstrate that relative rotation of the domains is a rate-limiting step of polymerization. Our structural analysis of interdomain interactions suggests that R- to T-state transition likely follows addition of a GTP-bound monomer to the filament through interaction of the preformed NTD. Hence, the addition of monomers to the NTD-exposed end of filament is slower in comparison to the C-terminal domain end, thus supporting the phenomenon of kinetic polarity in a single protofilament assembly.

## Introduction

FtsZ, a tubulin homolog, is one of the essential cytoskeletal proteins involved in bacterial cell division (Begg and Donachie, 1985; Nogales *et al*., 1998). To facilitate division, FtsZ localizes at the mid-cell forming a ring-like structure known as Z-ring (Bi and Lutkenhaus, 1991). The Z-ring tethers to the membrane via proteins like FtsA, an actin homolog, or ZipA, and recruits other proteins required for cell division (Ma *et al*., 1996; Hale and de Boer, 1997). This results in the formation of a multiprotein complex called divisome that constricts the cell, eventually leading to two daughter cells (Margolin, 2005). The precise mechanism of how ring constriction takes place and the role of FtsZ filament dynamics in the process is not fully understood. Most studies till date have focused on FtsZs from cell walled bacteria (de Boer *et al*., 1992; Mukherjee *et al*., 1993; Wang and Lutkenhaus, 1993; Anderson *et al*., 2004). Based on recent studies, cell wall synthesis is thought to be the primary force generator for constriction and the rate-limiting process in cytokinesis or septum closure (Coltharp and Xiao, 2017), (Bisson-Filho *et al*., 2017; Yang *et al*., 2017), (Monteiro *et al*., 2018; McCausland *et al*., 2021). However, how FtsZ drives cytokinesis in a cell-wall-less milieu is not understood. Hence, understanding the dynamics of FtsZ assembly from cell-wall less bacteria gains relevance.

FtsZ has a conserved domain architecture similar to tubulin, consisting of a GTP-binding N-terminal domain (NTD) and a C-terminal domain (CTD) connected via H7 helix (Löwe and Amos, 1998), followed by a flexible region that helps FtsZ to bind membrane binding proteins like ZipA or FtsA (Haney *et al*., 2001; Margolin, 2003; Cohan *et al*., 2020). FtsZ makes homopolymers comprising a single protofilament, in contrast to tubulin, which forms a heteropolymeric (consisting of alpha and beta tubulin) 13-stranded microtubule assembly. FtsZ polymerization is a cooperative process that requires a critical concentration of monomers for filament assembly (Mukherjee, 1998; Huecas and Andreu, 2003; Huecas *et al*., 2008; Surovtsev *et al*., 2008). In addition, FtsZ filaments exhibit treadmilling, as opposed to the typical dynamic instability observed in microtubules. A conformational switch of each monomer from its globular state (R-state; relaxed state favored by monomers in solution) to a filament compatible state (T-state; tense state adopted by monomers within a filament) favors cooperative assembly of FtsZ filaments (Huecas *et al*., 2008; Dajkovic *et al*., 2008; Wagstaff *et al*., 2017). This is a common feature shared across diverse tubulin family members that contribute to their cytomotive properties (Wagstaff *et al*., 2023). In the R-state, the inter-domain cleft between the NTD and the CTD is closed whereas in the T-state, it is open (**Figure 1A**; (Fujita *et al*., 2017; Wagstaff *et al*., 2017)). Other signature features of T-state conformation include a straightened orientation of the guanine ring and a downward movement of H7 helix (**Figure 1A**; (Wagstaff *et al*., 2017)). It has been hypothesized that the transition from R- to T-state within a monomeric subunit is thermodynamically less favorable due to the requirement of a positive free energy for opening of the inter-domain cleft (Corbin and Erickson, 2020). However, within protofilaments, the interactions between adjacent subunits provide the required energy for opening the cleft and favors R- to T-state transition to form a protofilament (Corbin and Erickson, 2020).

**Figure 1.**
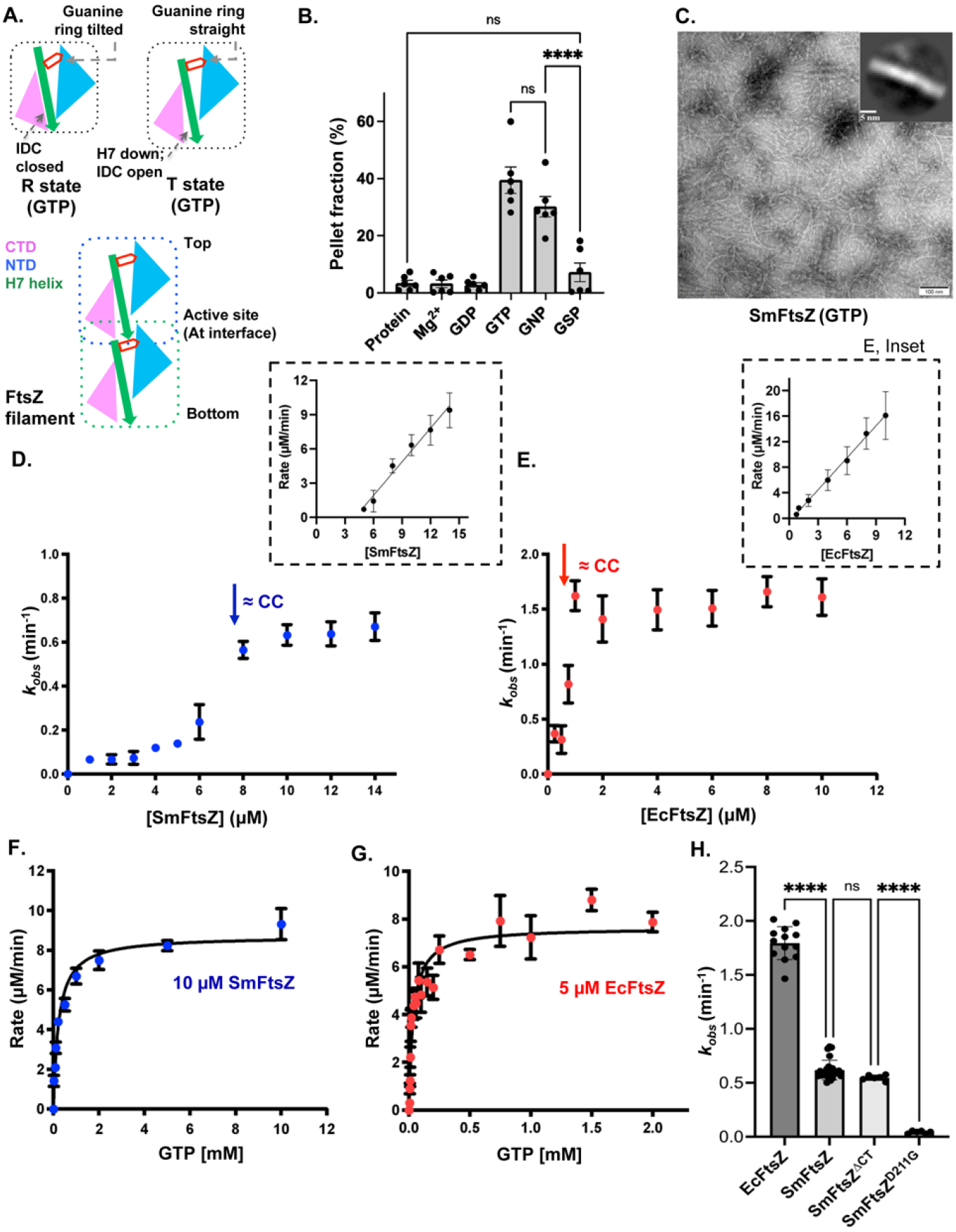
GTPase activity measurements highlight higher critical concentration and lower GTPase activity for SmFtsZ compared to EcFtsZ. **A)** Schematic showing FtsZ molecules in R- and T-states. The nucleotide hydrolysis occurs at the interface, and hence is a read-out for critical concentration of FtsZ. The NTD is in sky blue; the H7 helix and the T7 loop is denoted in green; the CTD is in pink; the pentagon denotes GTP. The various conventions of FtsZ filaments described in the text are indicated. **B)** Quantification for pelleting assay of SmFtsZ. Incubation of SmFtsZ (10 μM) in presence of 5 mM Mg^2+^; 3 mM GDP and 5 mM Mg^2+^; 3 mM GTP and 5 mM Mg^2+^; 3 mM GMPPNP (GNP) and 5 mM Mg^2+^; and 3 mM GTPγS (GSP) and 5 mM Mg^2+^, in pH 6.5. The amount of protein in the pellet provides an estimate of the filament fraction (a representative gel is shown in Figure S1C) (*N* = 2, *n* = 6). The error bar shows mean with SEM. **C)** Visualization of the SmFtsZ (10 µM) filaments obtained in presence of 2 mM GTP and 5 mM Mg^2+^, using transmission electron microscopy confirmed the presence of filaments. Scale bar represents 100 nm. The inset shows a 2D class average indicating the presence of single protofilaments of FtsZ and the scale bar represents 5 nm. **D)**, **E)** GTPase activity of SmFtsZ and EcFtsZ measured at increasing protein concentrations, respectively (SmFtsZ [WT; *N* = 3; *n* = 4], EcFtsZ [WT; *N* = 3; *n* = 5]. Insets show the plot of rate vs concentration of proteins, x-intercept for SmFtsZ = 4.1 µM and that of EcFtsZ = 0.52 µM. **F)**, **G)** GTPase activity of 10 µM SmFtsZ and 5 µM EcFtsZ, respectively, measured at increasing concentrations of GTP ranging from 0 to 10,000 µM (blue) for SmFtsZ and 0 to 2000 µM (red) for EcFtsZ. The curve was fitted with the Michaelis-Menten equation, V_0_ = V_max_ ([S]/([S] + K_M_). (*N* = 3; *n*= 3). **H)** Comparison of GTPase activity (*k_obs_* defined as relative specific activity (min^-1^)) of FtsZs. SmFtsZ^D211G^, a GTPase dead mutant, was taken as negative control. The experiments were done with saturating concentrations of GTP (2.5 mM for SmFtsZ and 1 mM for EcFtsZ). (EcFtsZ [*N* = 4; *n*= 13], SmFtsZ [*N* = 3; *n* = 25], SmFtsZ^ΔCT^ [*N* = 2; *n* = 6], and SmFtsZ^D211G^ [*N* = 1; *n* = 6]; The protein concentration used was 10 µM. The mean with SEM and 95% confidence intervals is shown in the long and short horizontal lines in panel H, one-way ANOVA; \*\*\*\**p* < 0.0001; ns, non-significant. The estimated *k_obs_* values are tabulated in **Table S2**. *N* = number of independent purification batches and *n* = number of repeats.

A crucial aspect of the dynamics of the FtsZ filament is its kinetic polarity, which is interestingly the opposite of what is known about microtubules and was inferred from mutational analysis (Du *et al*., 2018). Microtubules elongate from the end at which the N-terminal domain of beta-tubulin is exposed (referred to as the top end hereafter), whereas FtsZ elongates from the end with an exposed C-terminal domain (referred to as the bottom end hereafter) (Du *et al*., 2018; Knossow *et al*., 2020). However, the structural basis for the directionality of polymerization is not completely understood, though both the homologs undergo a similar conformational change during the switch from R- to T-state (Wagstaff, et al, 2023).

In this study, we characterized FtsZ from *Spiroplasma melliferum* (SmFtsZ), which is a cell wall-less bacteria, and compared it with the relatively well-characterized FtsZ from *Escherichia coli* (EcFtsZ), a cell walled bacterium. *Spiroplasmas* are cell-wall-deficient bacteria that belong to the Mollicutes class and are distinguished by their pleomorphic helical cell structure (Razin *et al*., 1973). Only 5 (*mraZ*, *mraW*, *ftsA*, *ftsZ*, and *sepF*) of the over 15 genes coded by the division and cell wall (*dcw*) cluster in cell-walled bacteria are present in most Mollicutes (Zhao *et al*., 2004; Harne *et al*., 2020). These five proteins make up the necessary machinery for cell division in bacteria lacking a cell wall, given the lower genome sizes of these bacteria. For *Spiroplasma*, two cell division strategies have been proposed. Based on the observation of the growth of *S. citri*, it was inferred that *Spiroplasma* cells split by one or more constrictions along the short axis of the cell and develop at one of the two poles by insertion of additional membrane components (Garnier *et al*., 1981; Garnier *et al*., 1984). Another proposed mode of division is the longitudinal fission or the Y shaped fission. The immunogold labeling by anti-FtsZ confirmed the presence of FtsZ in the branching area of the Y shaped cell, indicating that the division is dependent on FtsZ (Ramond *et al*., 2016). We chose SmFtsZ from cell wall-less bacteria as a model to study fundamental features of FtsZ hydrolysis and filament dynamics in ring constriction and cytokinesis, independent of cell wall synthesis.

Biochemical analysis of SmFtsZ revealed a lower GTPase activity and higher critical concentration of filament formation than EcFtsZ. SmFtsZ formed ring-like structures in *E. coli* indicating its polymerization ability. Crystal structures showed the presence of a domain-swapped dimer, in which the swapped domains organize to mimic the globular structure of FtsZ monomers. The crystal structures of SmFtsZ and comparison with other FtsZ structures in different nucleotide states suggested that the gamma phosphate of GTP analogs induces conformational changes in the T3 loop that might trigger polymerization. Our *in vitro* and *in vivo* characterization of the inter-domain cleft mutants of SmFtsZ and EcFtsZ support the relevance of Phe 224 (SmFtsZ numbering), a residue in the cleft between NTD and CTD, in driving the facile conversion to the T-state. Based on the structures of FtsZ conformational states, biochemistry and *in vivo* experiments, we propose that the cleft opening is a rate limiting step of polymerization. The conformational switch from R to T-state supports a preferred addition from bottom end of the filament, and thus provides a basis for the kinetic polarity of FtsZ.

## Results

### SmFtsZ has higher critical concentration and lower GTPase activity than EcFtsZ *in vitro*

We purified SmFtsZ (**Figure S1A, B**) and performed sedimentation assay in order to assess the ability of SmFtsZ to polymerize. SmFtsZ was observed in the pellet fraction (representing polymerized fraction of FtsZ) specifically in the presence of GTP or GMPPNP (GNP) and MgCl_2_ and not upon addition of GDP or GTPγS (GSP) and MgCl_2_ (GMPPNP: guanosine 5’-[β, γ-imido] triphosphate, GTPγS: guanosine 5’-O-[gamma-thio] triphosphate; non-hydrolyzable analogs of GTP (**Figure 1B; S1C**). Further, observation of SmFtsZ filaments in presence of GTP and Mg^2+^ using electron microscopy and their 2D class averages (**Figure 1C**) led us to conclude that SmFtsZ formed single filaments in the presence of GTP similar to other characterized bacterial FtsZs (Wagstaff *et al*., 2017).

As the GTPase active site is at the interface between two monomers in the FtsZ filament (**Figure 1A**), filament assembly is a requirement for GTP hydrolysis. Hence, we used the rate of GTP hydrolysis as a proxy to estimate critical concentration (CC) of filament formation. An approximate estimate of CC will be the concentration above which *k_obs_* (defined as the amount of GTP hydrolysis per unit concentration of the enzyme per unit time) reaches a constant value. For SmFtsZ, we noticed that after a concentration of 8 μM, a constant activity of 0.62 ± 0.02 min^-1^ (average of *k_obs_* of the protein concentrations above estimated CC) was observed (**Figure 1D**). For EcFtsZ, a constant *k_obs_* of 1.54 ± 0.06 min^-1^ (average of *k_obs_* of the protein concentrations above estimated CC) was observed above a lower value of approximately 1 μM (**Figure 1E**). The x-intercept of rate vs protein concentration provided an alternative estimate of the critical concentrations with 4.1 µM and 0.52 µM for SmFtsZ and EcFtsZ, respectively (**Figure 1D, E, insets**).

We then measured the specific activity of these two proteins at a saturating concentration of GTP, the GTP concentration being fixed based on the Michaelis-Menten plot (**Figure 1F, G**). The specific activities (*k_cat_*) were estimated as 0.61 ± 0.02 min^-1^ and 1.79 ± 0.04 min^-1^ for SmFtsZ and EcFtsZ, respectively, consistent with the values above, indicating an approximately two- to three-fold lower activity for SmFtsZ (**Figure 1H**). Activity measurements of the active site mutant SmFtsZ^D211G^ showed that the mutation abrogated GTPase activity and SmFtsZ^ΔCT^, a truncated construct of SmFtsZ without the C-terminal flexible linker, showed similar activity as the wild type (**Figure 1H**). Importantly, the Michaelis-Menten plot also revealed that the *K_m_* of SmFtsZ (234 μM) was 7 times higher than that of EcFtsZ (34 μM) (**Table S1**). Hence, the catalytic efficiency for SmFtsZ was 12 times lower than that of EcFtsZ (**Table S1**).

Thus, our biochemical data demonstrated that SmFtsZ possesses a higher critical concentration for filament formation with a lower GTPase activity compared to EcFtsZ (**Tables S1, S2**) under similar *in vitro* conditions.

### SmFtsZ forms ring-like structures in *E. coli*

We next sought to know whether SmFtsZ can polymerize into rings *in vivo*. We targeted SmFtsZ directly to *E. coli* membranes by generating clones for SmFtsZ chimeric construct in which the core of SmFtsZ was tagged with a translational fusion of mNeonGreen (mNG) with the membrane targeting sequence (mts) of EcMinD at its C-terminus to check whether it can form rings in *E. coli in vivo* (**Figure 2A**). We cloned SmFtsZ core (1 - 314 amino acids) region into pJSB100 plasmid replacing EcFtsZ core (1 – 366 amino acids) and having mNeongreen tag with the MinD membrane targeting sequence (mts) at its C-terminus. This SmFtsZ chimeric construct was expressed in the *E. coli* strain BW27783 (CCD161), which localized as scattered rings or spiral polymers attached to the membrane throughout the cell including cell poles (EcFtsZ as control; **Figure 2B, C).**

**Figure 2.**
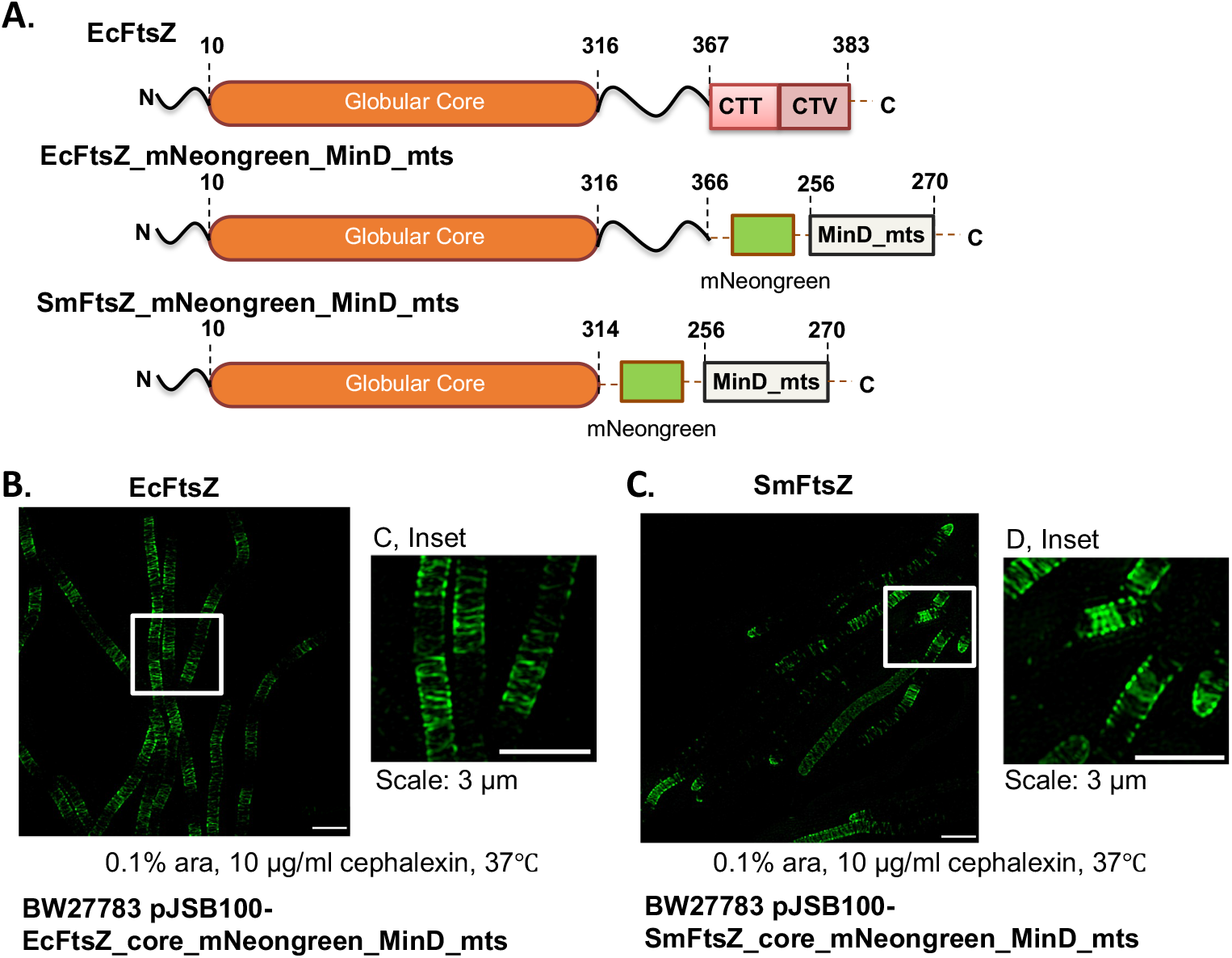
SmFtsZ polymerizes to form rings in *E. coli in vivo*. **A)** Schematic of the constructs used in the experiment namely EcFtsZ, EcFtsZ_mNeongreen_MinD_mts, and SmFtsZ_mNeongreen_MinD_mts. The residue numbering denotes the extent of corresponding sequences in the individual proteins. **B)**, **C)** Phenotypic localization of SmFtsZ when expressed in *E. coli* BW27783 (CCD161) strain carrying pJSB100 plasmid with EcFtsZ (1-366) and SmFtsZ (1-316) core domain tagged with mNeonGreen at its C-terminus followed by EcMinD_mts, respectively were imaged using three-dimensional structured illumination microscopy (3D-SIM). Maximal intensity projection images are shown.

This demonstrates that SmFtsZ indeed possesses the features of FtsZ that enables it to polymerize in vivo and form ring-like structures in *E. coli*.

### Crystal structures of SmFtsZ^ΔCT^ captured as domain swapped dimers

To obtain mechanistic insights on SmFtsZ activity, we purified and crystallized SmFtsZ^ΔCT^ (a truncated construct of SmFtsZ without the C-terminal flexible linker; **Figure S1A, B**), without and with addition of GMPPNP. Crystal structures of both GDP-bound SmFtsZ^ΔCT^ (GDP intrinsically bound to the protein during purification; **Figure S1D, S1E**) (PDB ID: 7YSZ) and GMPPNP-bound SmFtsZ^ΔCT^ (**Figure S2A, B**) (PDB ID: 7YOP) were obtained in the same space group and unit cell, implying similar crystal packing (**Table S3**). The structure 7YSZ has two molecules in the asymmetric unit and both the chains are bound to GDP (**Figure S2A**). In contrast, the structure 7YOP, which too, has two molecules in the asymmetric unit, has GDP bound to chain A and GMPPNP bound to chain B (**Figure S2B**). Interestingly, the two molecules in the asymmetric unit exist as a domain-swapped dimer, wherein the NTD of one chain is packed against the CTD of the other chain, which together resemble the globular structure of FtsZ (**Figure 3A**). In the extended conformation of one molecule of the domain-swapped dimer (**Figure 3B**), the NTD and the CTD were separated at the T6 loop which is the start of the H7 helix. A clear and continuous electron density in the T6 loop region was observed in this region for chain B (**Figure 3B**, **inset**), confirming the domain-swapped conformation of SmFtsZ^ΔCT^ in the crystal structure. For chain A, the corresponding loop was disordered and hence the residue stretch of 170–185 has not been modelled (**Figure 3B**, **inset**). The relative orientations of the NTD and CTD in the extended conformation of each chain (A and B) are different (**Figure 3B**).

**Figure 3.**
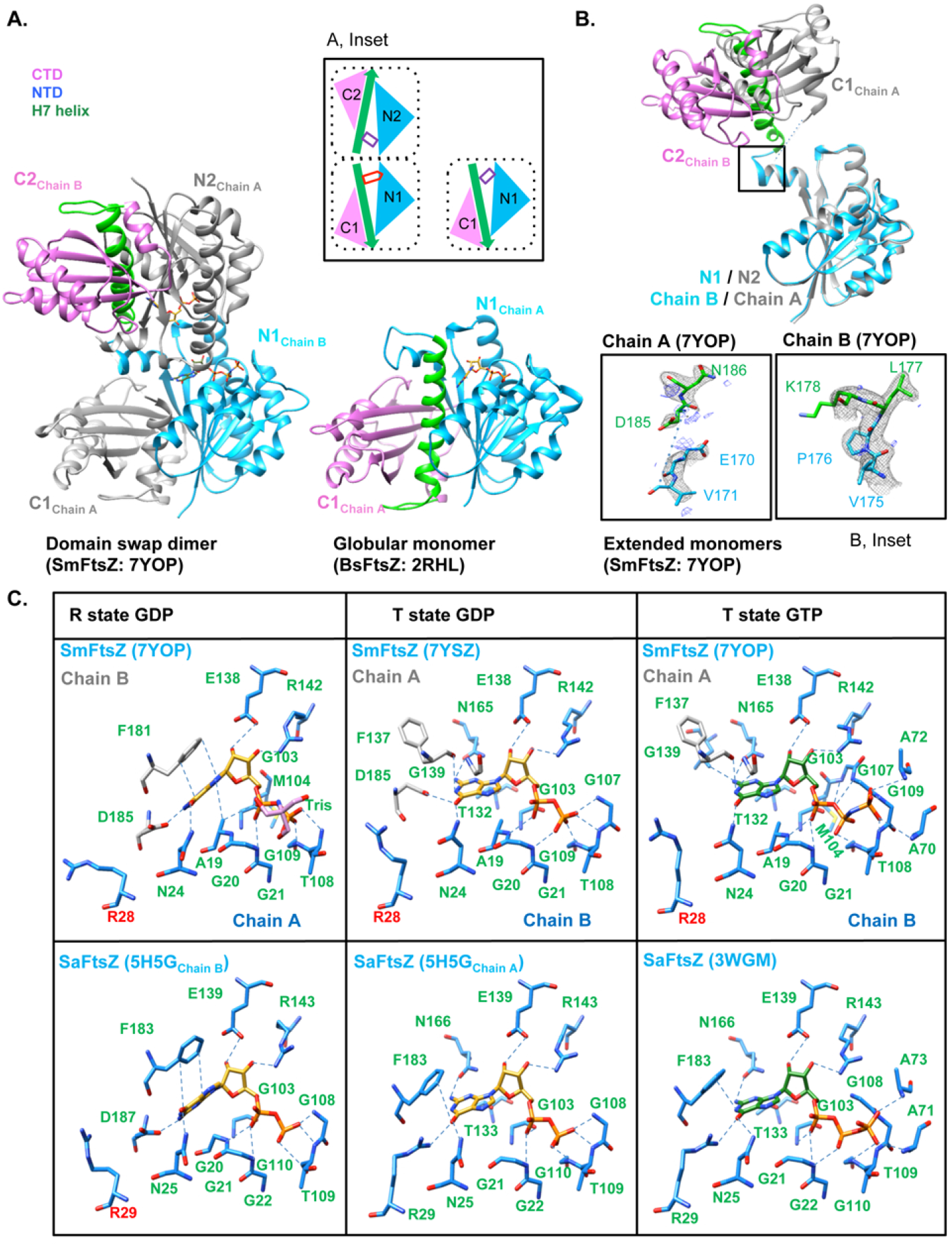
SmFtsZ crystallized as a domain swap dimer. **A)** Overall structure of SmFtsZ, along with the schematic containing both the chains forming domain-swap dimers. The domains and the chains are denoted with the respective colors as in the schematic (PDB ID: 7YOP). Overall structure of globular FtsZ from *B. subtilis* (PDB ID: 2RHL). The inset shows the schematic of the structure, where the domains are denoted as: NTD (blue), H7 helix and T7 loop (green), and CTD (pink). Purple rectangle denotes GDP and the red pentagon denotes GTP. **B)** Relative orientations of the CTD in the extended conformation of each chain, fixing C-alpha of NTD of both the chains. Electron density map of the region shown in the inset where the two domains separate at T6 loop (map *F_o_* – *F_c_* shown at 2.5 σ, blue and 2*F_o_* – *F_c_* at 1.2 σ, black). **C)** Nucleotide conformation of SmFtsZ along with SaFtsZ in R-state in GDP (5H5G, chain B), SaFtsZ in T state in GDP (5H5G, chain A), GTP (3WGM) are shown (GDP - golden, GTP - forest green).

Despite the presence of a domain-swapped dimer in the crystal structure, the SEC-MALS profile indicates predominantly a monomeric nature for SmFtsZ^ΔCT^ in solution at lower concentrations of the protein (**Figure S2C**). At a higher concentration, the SEC profile shows two peaks - a major peak corresponding to a monomer form and a minor peak corresponding to a dimeric form (**Figure S2D, E**). Since the polymerization interface is occluded in the domain-swapped conformation, the conformation cannot polymerize, which might have facilitated preferential crystallization of the domain-swapped dimer. Though no nucleotide was added during crystallization, GDP was bound in the nucleotide pocket in both chains of one of the structures (PDB ID: 7YSZ) (**Figure S2A**). It is likely that the bound GDP was from *E. coli* cells during expression of SmFtsZ^ΔCT^ (seen in HPLC of denatured protein extract of purified protein, **Figure S1E**). Upon quantification of the HPLC data, it was seen that the ratio of SmFtsZ to GDP was approximately 1:1, indicating that all molecules of SmFtsZ are bound to GDP (**Figure S1D, E**). In the second structure (PDB ID: 7YOP) where GMPPNP was added prior to crystallization, GDP and GMPPNP were observed in the nucleotide-binding pockets of chain A and chain B, respectively (**Figure S2B**).

To further investigate the structural details of the unique conformations of SmFtsZ, we compared the structures with R-state GDP bound (PDB ID: 5H5G, chain B), T-state GDP bound (PDB ID: 5H5G, chain A), and T-state GTP bound (PDB ID: 3WGM) conformations of SaFtsZ (*Staphylococcus aureus*), respectively. In the structures of chain A of PDB 7YSZ and PDB 7YOP, the orientation of the guanine ring of GDP matched that in the R-state of FtsZ, while in chain B of both 7YSZ (GDP bound) and 7YOP (GMPPNP bound), the orientation of the nucleotide base corresponded to that in the T-state (**Figure 3C**). The angle between the planes of the guanine rings in the R and T state for all the available crystal structures of FtsZ (**Table S4**) confirmed that the conformational state of the nucleotide in SmFtsZ corresponded to R-state and T-state in chain A and B, respectively (**Figure S2F**).

Typically, a phenylalanine residue from H7 helix (Phe183) stacks against the nucleotide base to stabilize the T-state conformation in the FtsZ filament. The guanine ring orientation is held in the T-state position due to crystal packing interactions with the adjacent monomer in the domain swapped SmFtsZ structures. An analysis of the relative positions of the NTD, CTD and H7 helix showed that the NTD, CTD and H7 helix positions in both the chains corresponded to the R-state conformation for both the structures-7YSZ and 7YOP (**Figure S2G**), despite the nucleotide orientation in NTD of chain B resembling the T-state orientation. The nucleotide orientation is decoupled from the orientation of the Phe183 of the H7 helix in SmFtsZ structures due to the domain swap, while it is usually correlated with the H7 helix position in globular structures of FtsZ.

### Gamma-phosphate of the nucleotide stabilizes the conformation of the T3 loop compatible with polymerization

To analyze the conformational changes in the GMPPNP-bound state in comparison to the GDP-bound state, the NTD of GDP-(chain A and B of 7YSZ, chain A of 7YOP) and GMPPNP-bound (chain B of 7YOP) structures were superposed. A clear distinction was seen in the T3 loop, which surrounds the gamma phosphate and is also involved in interaction with the subunit above in the filament (Yoshizawa *et al*., 2020). We found that in the GDP-bound state of chain A (PDB ID: 7YSZ and 7YOP), the T3 loop was ordered with the carbonyl group of Gly71 of T3 loop facing towards the nucleotide phosphate. A water-mediated interaction between carbonyl oxygen of Gly71 and with the beta phosphate of GDP was observed in chain A of both the structures (**Figure 4A, B**).

**Figure 4.**
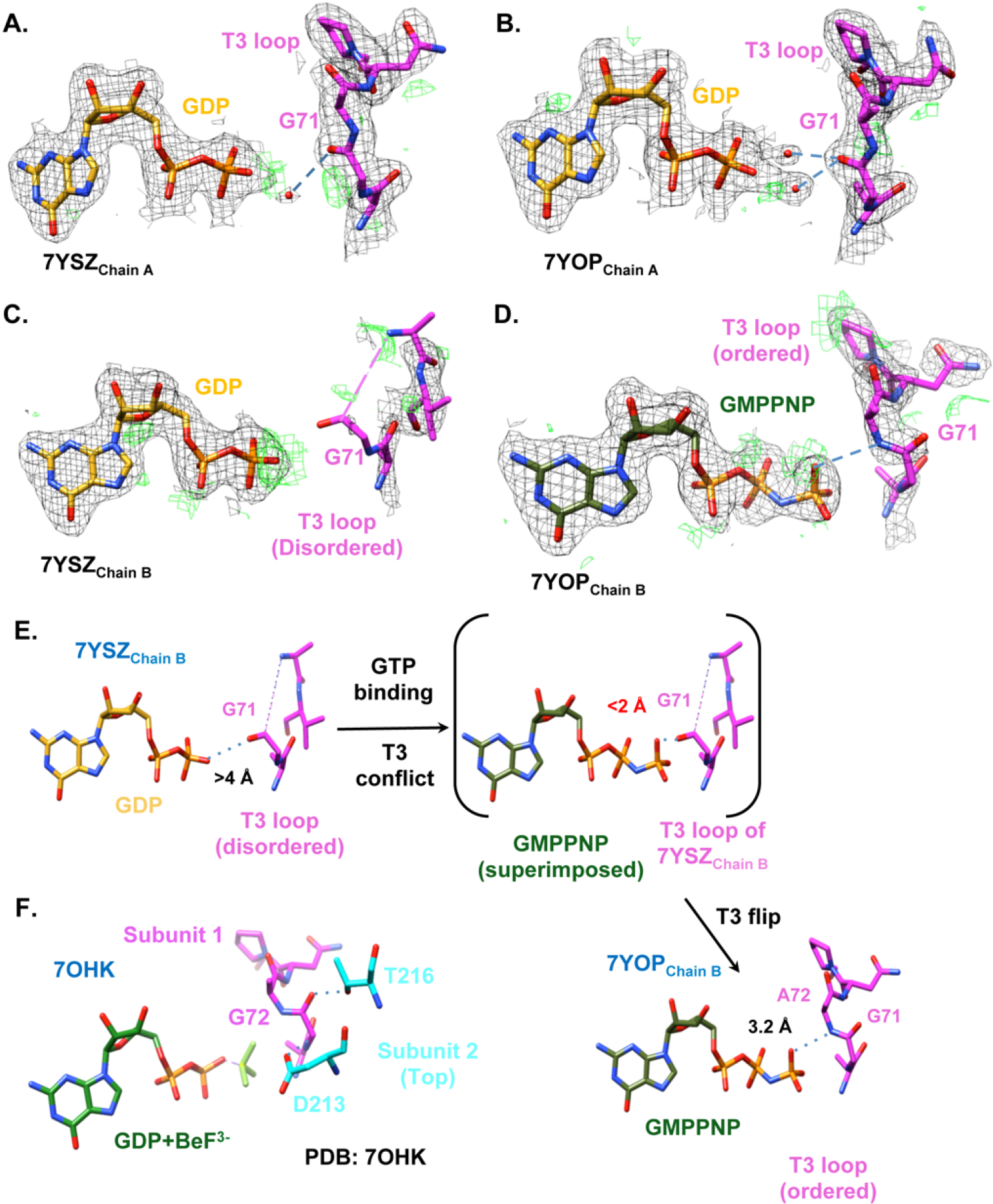
Filament compatible conformation of T3 loop is stabilized by gamma-phosphate of the nucleotide. **A)**, **B)** Electron density map for the orientation of T3 loop in R-state GDP-bound structures of SmFtsZ (PDB ID: 7YSZ, chain A; PDB ID: 7YOP, chain A, respectively). **C)**, **D)** Electron density map for the orientation of T3 loop in T-state GDP- and GMPPNP-bound structures of SmFtsZ (PDB ID: 7YSZ, chain B; PDB ID: 7YOP, chain B, respectively). Map *F_o_* – *F_c_* shown at 2.5 σ (green) and 2*F_o_* – *F_c_* at 1.2 σ (black). **E)** Comparison of SmFtsZ structure bound to GDP (chain B, PDB ID: 7YSZ) and SmFtsZ bound to GMPPNP (chain B, PDB ID: 7YOP). The carboxyl group of Gly71 of T3 loop faces towards the nucleotide pocket. The loop is disordered in the presence of GDP in the T-state. GMPPNP binding induces steric hindrance with the Gly71 peptide of T3 loop due to which the peptide flips. **F)** Filament structure of FtsZ from *Staphylococcus aureus* (PDB ID: 7OHK). The Gly72 peptide (T3 loop) flips out (subunit 1) and interacts with Asp213 and Thr216 of the upper subunit (subunit 2).

In the GDP bound chain B (7YSZ), the loop was only partially modeled due to poor electron density. In the GMPPNP-bound chain B (7YOP), the peptide of Gly71 of the loop was modeled in a flipped-out conformation, away from the nucleotide (**Figure 4C, D**), an essential change for accommodating the gamma phosphate without steric clash with the carbonyl oxygen of Gly71 (**Figure 4E**). The flipped conformation of the peptide matches with that of the filament interacting conformation of T3 loop (as observed in 7OHK; **Figure 4F**) with the gamma phosphate conformation of GMPPNP matching that of GTP. Based on these observations, we propose that the flipping of Gly71 and stabilization of the T3 loop is a key structural feature for the GTP-triggered polymerization of FtsZ (**Figure 4F**). Indeed, in all the structures of FtsZ, Gly71 is always flipped out in the presence of GTP or its analogs. In the presence of GDP, the crystal packing stabilizes the T3 loop, resulting in some instances where the T3 loop is flipped even in the GDP bound structures (**Figure S2H**). The absence of an adjacent subunit that satisfies the contacts from a polymerization interface possibly resulted in capturing the partial disorder of the T3 loop of the two T-state conformations in chain B of 7YOP and 7YSZ, respectively.

These results suggest that the Gly71 peptide of T3 loop is flipped out in presence of nucleotide with gamma phosphate, which in turn is involved in filament interface interaction.

### A flexible residue in the inter-domain cleft facilitates state transition in FtsZ

Based on our biochemical characterization and observations from the crystal structure, SmFtsZ appeared to have a higher critical concentration and a tendency to form an extended structure, a state achieved probably during the transition from R- to T-state. A key step for the R-state to T-state transition, which determines the critical concentration and kinetics of polymerization, is the relative rotation between NTD and CTD leading to the opening of the cleft between them. We identified the interacting residues across the cleft and analyzed their conservation across FtsZ sequences from different organisms (**Figure S3A**). A notable difference was the observation of phenylalanine (Phe224) in strand S7 of SmFtsZ, instead of methionine at the equivalent position in most other FtsZs (including EcFtsZ or *Staphylococcus aureus* FtsZ; SaFtsZ) (**Figure 5A**). The conformations of this residue in the SmFtsZ^ΔCT^ structures are shown (**Figure 5A insets**). We hypothesized that the greater flexibility of methionine in comparison to phenylalanine might facilitate the transition from one state to another (e.g., in EcFtsZ).

**Figure 5.**
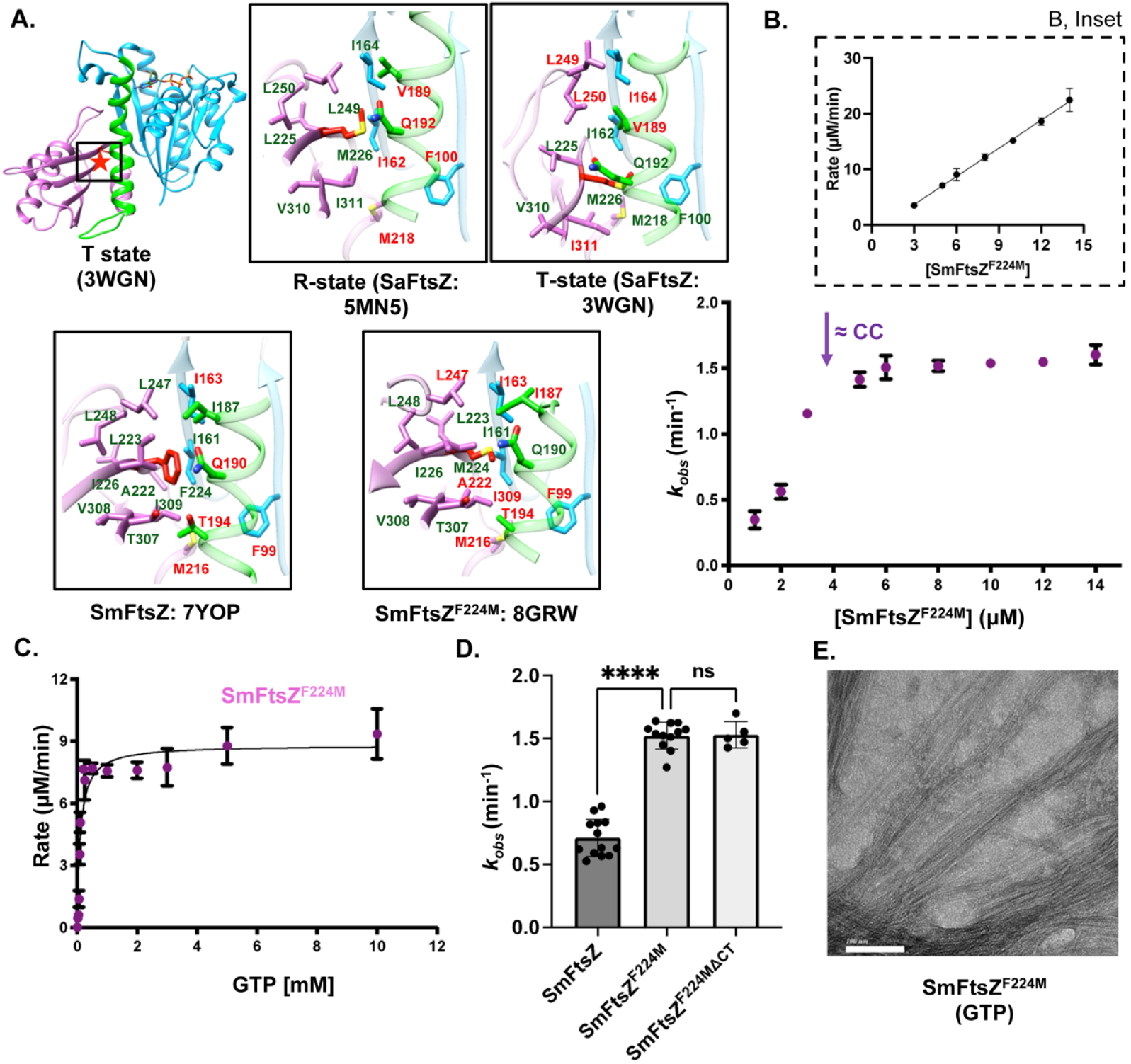
Phe224 plays a role in the ease of transition from R- to T-states and is a hyperactive mutant of SmFtsZ. **A)** Overall structure of FtsZ (PDB ID: 3WGN), the boxes highlight the position of the methionine, the residue of interest. The insets show the position of Met226 in SaFtsZ and SmFtsZ in S7 strand comparing R-state (SaFtsZ; PDB ID: 5MN5), T-state (SaFtsZ; PDB ID: 3WGN), chain A of SmFtsZ (PDB ID: 7YOP), and chain A of SmFtsZ^F224M^ (PDB ID: 8GRW) highlighting the movement of the CTD. **B)** GTPase activity of SmFtsZ^F224M^ was measured with different concentrations of the protein ranging from 0 to 14 µM, [*N* = 2; *n* = 4]. The inset shows the plot of rate vs concentration of protein; x-intercept = 0.84 µM. **C)** GTPase activity of 5 µM SmFtsZ^F224M^ was measured with different concentrations of GTP ranging from 0 to 10,000 µM (purple). The curve was fitted with the Michaelis-Menten equation, V_0_ = V_max_ ([S]/([S] + K_M_), [*N* = 2; *n* = 4]. **D)** GTPase activity comparison of SmFtsZ^F224M^ mutant with the wild type construct (SmFtsZ^F224M^ [*N* = 2; *n* = 12], SmFtsZ^F224M^^ΔCT^ [*N* = 2; *n* = 5] and SmFtsZ [*N* = 3; *n* = 15]). The protein concentration used was 10 µM. The mean with SEM and 95% confidence intervals is shown in the long and short horizontal lines in panel D, one-way ANOVA; \*\*\*\**p* < 0.0001; ns, non-significant. **E)** Visualization of the SmFtsZ^F224M^ (10 µM) filaments obtained in presence of 2 mM GTP and 5 mM Mg^2+^, using transmission electron microscopy confirmed the presence of filaments. Scale bar represents 100 nm.

To test our hypothesis, we mutated Phe224 to methionine in SmFtsZ^ΔCT^ (SmFtsZ^F224M,ΔCT^), purified the mutant protein (**Figure S3B, C**) and performed biochemical assays. To find if the mutation confers any change in the process of polymerization (by comparison of critical concentrations), GTPase assay with different concentrations of SmFtsZ^F224M^ was performed. It was observed that the *k_obs_* of 1.47 ± 0.03 remained constant above a protein concentration of approximately 3 µM for SmFtsZ^F224M^ (**Figure 5B**). The x-intercept of rate vs protein concentration provided an alternative estimate of the critical concentration, which is 0.84 µM. This indicated that the critical concentration of the mutant was lower than the wild type (4.1 µM). We observed from the Michaelis-Menten plot (**Figure 5C**) that the catalytic efficiency of this mutant was four times higher (**Table S1**) and the GTPase activity (*k_obs_*) was approximately three times higher than that of the wild type (**Figure 5D, Table S2**). Filaments of SmFtsZ^F224M^ in presence of GTP were observed under TEM (**Figure 5E**). We also obtained the structure of SmFtsZ^F224M,ΔCT^ in presence of GDP (PDB ID: 8GRW, **Table S3**) but the mutant could not be crystalized with GMPPNP, possibly due to efficient polymerization being inhibitory to crystallization (**Figure S3D**). Although the GDP-bound SmFtsZ^F224M,ΔCT^ structure was a domain-swapped dimer similar to that of the wild type structure, the methionine at position 224 interacted with fewer residues as compared to that of phenylalanine in wild type SmFtsZ (**Figure 5A, insets**). Given the similar crystal packing, the nucleotide conformations in the two chains were also similar to that of the wild type structure.

Thus, based on the CC and the GTPase activity measurements, the cleft mutant, SmFtsZ^F224M^, demonstrated that a smaller and more flexible residue at position 224 (residue numbering as in SmFtsZ) in the inter-domain cleft might lead to a quicker transition from R- to T-state.

### A rigid phenylalanine instead of flexible methionine in the inter-domain cleft in EcFtsZ affects *E. coli* cell division

EcFtsZ and many other bacterial FtsZs have a methionine at the position equivalent to SmFtsZ-Phe224. We, hence, sought to test if a phenylalanine at this position in EcFtsZ would affect FtsZ function *in vivo*. As described above, to test if EcFtsZ^M225F^ affects cell division in absence of EcFtsZ, we utilized the *E. coli* strain JKD7-1 carrying the plasmid pKD3 (Stricker and Erickson, 2003). Only EcFtsZ or only mutants of EcFtsZ that can support division in the absence of FtsZ^WT^ would give rise to viable colonies on plates containing arabinose at 42 °C.

We therefore first tested, using spotting assay, if EcFtsZ^M225F^ mutant supported growth of JKD7-1 strain at 42 °C, i.e., in the absence of any endogenous EcFtsZ protein. EcFtsZ served as the positive control and the empty vector pJSB2 as the negative control. While all of them grew well at 30 °C in the presence of 1% glucose (added as a repressor to prevent any expression from the arabinose promoter *araBAD*, in pJSB2 derived plasmids), no colony formation was observed with the vector alone (pJSB2) at 42 °C as expected. Both EcFtsZ and EcFtsZ^M225F^ formed colonies on plates with 0.2 % arabinose under permissive temperature of 30 °C, which also expressed the wild type FtsZ through the pKD3 plasmid as well as the restrictive temperature of 42 °C (**Figure 6A**). However, a microscopic examination of the EcFtsZ^M225F^ mutant at the restrictive temperature of 42 °C in liquid cultures revealed aberrant cell division with many filamentous cells (**Figure 6B**), indicating that the mutant was not fully functional. An analysis of their cell lengths showed that the cells expressing EcFtsZ^M225F^ exhibited filamentous morphology and were much longer than those expressing EcFtsZ. The average cell lengths of the wildtype and M225F mutant were found to be 4.58 µm ± 0.09 (number of cells counted were at least 218, N = 3) and 12.45 µm ± 4.47 (number of cells counted were at least 100, N = 3) respectively (**Figure 6C**).

**Figure 6.**
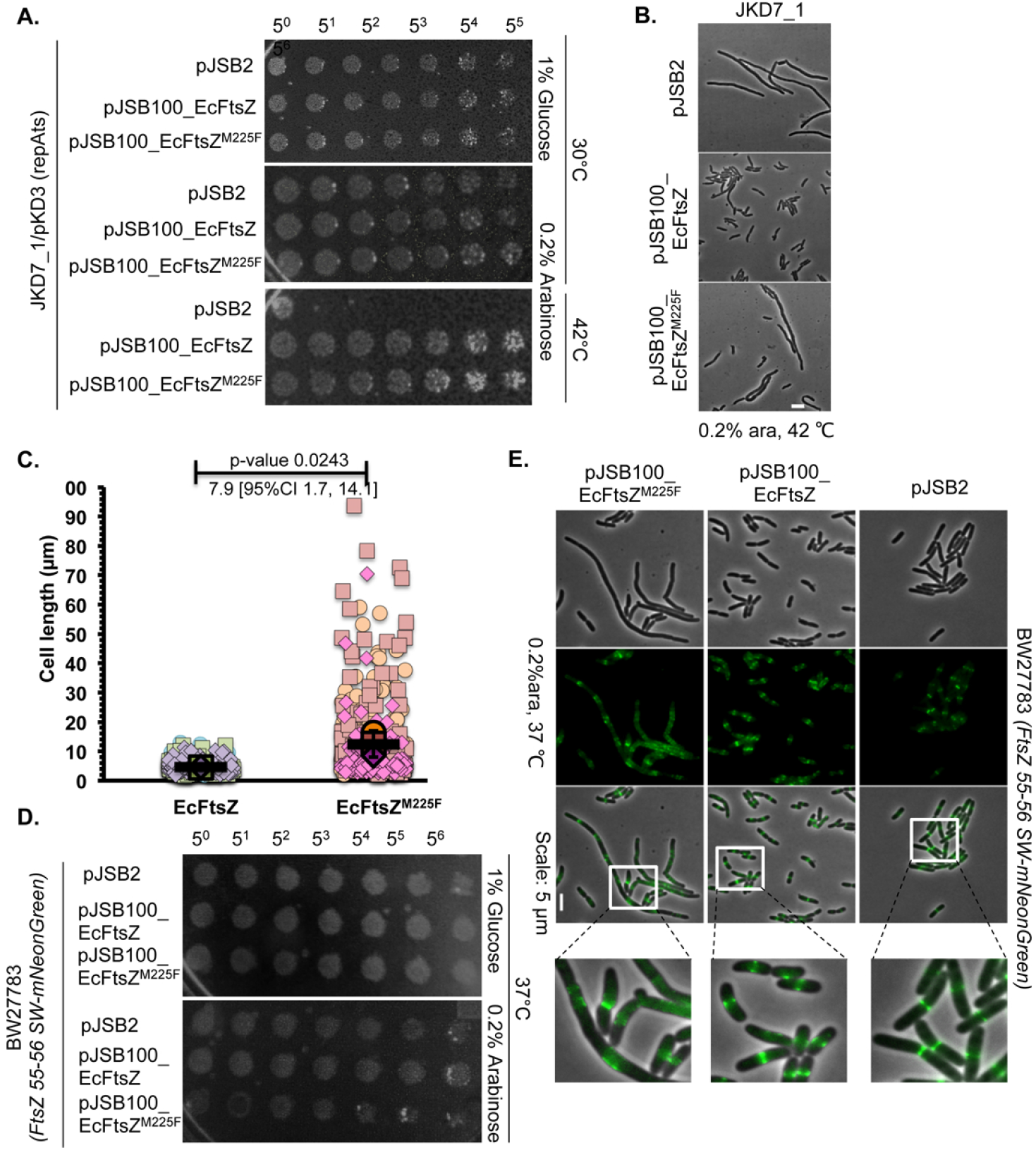
M225F mutation in EcFtsZ impairs *E. coli* growth and cell division. **A)** JKD7_1/ pKD3 strain carrying pJSB2 (vector) or pJSB100 (EcFtsZ) or pJSB100-EcFtsZ^M225F^ plasmids were spotted on the repressible plate (1 % glucose) or inducible plate (0.2 % L-arabinose). The plates containing L-arabinose were incubated at either permissible (30 °C) or non-permissible (42 °C) temperatures. Each subsequent spot was five times serially diluted. **B)** The cell morphology of the JKD7_1 strain carrying pJSB2 (vector) or pJSB100 (EcFtsZ) or pJSB100-EcFtsZ^M225F^ plasmids were observed by phase-contrast imaging after 6 hours of growth at 42 °C, when the pKD3 plasmid is lost and expression of FtsZ is solely from the pJSB plasmids. **C)** The wildtype EcFtsZ expressing JKD7_1 strain had an average cell length of 4.58 µm ± 0.09 (number cells counted were at least 218, N=3) and the cells expressing mutant EcFtsZ^M225F^ were 12.45 µm ± 4.47 (number of cells counted were at least 100, N=3) long. The quantitative data are shown as super plots and P-values and effect size are indicated above the plots. **D)** CCD288 (BW27783_ FtsZ 55-56-mNeonGreen-SW) strains with pJSB100 plasmids spotted on the repressible and inducible plates at 37 °C with five times of serial dilution. **E)** FtsZ rings were imaged in CCD288 strain (BW27783_FtsZ 55-56-mNeonGreen-SW) carrying empty vector (pJSB2) or ectopically expressing EcFtsZ or EcFtsZ^M225F^ by epifluorescence microscopy. Ectopic expression of EcFtsZ^M225F^ resulted in aberrant FtsZ ring compaction and cell division. Scale bar = 5 µm. Statistical significance was assessed by paired two-tailed Student’s *t* test. P values were calculated using Microsoft Excel formula T-TEST. The error bars shown are inferential and represent 95% confidence interval (CI).

These results suggested that although EcFtsZ^M225F^ mutant supported colony formation, the mutation affected cell division in *E. coli* as observed from the filamentous phenotype of the cells.

### Imperfect ring assembly accompanies defective cell division

We next sought to assess the effect of expression of EcFtsZ^M225F^ mutant on FtsZ ring assembly. In particular, we hypothesized that if the change to phenylalanine slowed down the state of transition, the ectopic expression of EcFtsZ^M225F^ in *E. coli* cells carrying the chromosomal copy of FtsZ, might affect the cell growth and FtsZ ring assembly. In order to monitor FtsZ ring assembly, we utilized a strain of *E. coli* (CCD288; a BW27783 derivative), wherein the wildtype chromosomally encoded FtsZ is fluorescently tagged (FtsZ_55-56_-mNeonGreen-SW) and is functional (Moore *et al*., 2016; Yang *et al*., 2017). We expressed untagged version of either EcFtsZ or EcFtsZ^M225F^ from the arabinose inducible promoter of pJSB2 derived plasmid and assessed their effect on the growth of the *E. coli* strain and the FtsZ rings assembled by FtsZ_55-56_-SW-mNeonGreen at 37 °C. Expression of EcFtsZ^M225F^ mildly affected the growth of *E. coli* as compared to those expressing EcFtsZ (**Figure 6D, S4A**). Further, imaging and cell length calculations showed that the strain expressing EcFtsZ^M225F^ was almost 3-fold longer [average cell length of 11.73 µm ± 1.56 (number of cells counted were at least 226, N = 3)] than those expressing EcFtsZ [average length of 4.09 ± 0.81 µm (number of cells counted were at least 166, N = 3)]. These results further confirmed the division defects due to the expression of EcFtsZ^M225F^ (**Figure S4B**).

To test if the expression of EcFtsZ^M225F^ perturbed FtsZ ring assembly, we visualized the FtsZ rings assembled by the chromosomally encoded functional FtsZ_55-56_-SW-mNeonGreen fusion protein under similar conditions. The untagged EcFtsZ^M225F^ was expressed from the plasmid (pJSB2*-EcFtsZ^M225F^)* under the control of arabinose promoter and induced with the addition of 0.2% arabinose at 37 °C. As a control, the untagged EcFtsZ was expressed in a similar manner. Western blotting was performed to ascertain that both wildtype and the EcFtsZ^M225F^ were expressed at similar levels (**Figure S4C, D**). In cells expressing EcFtsZ, the Z-rings (visualized due to endogenous mNeonGreen tagged EcFtsZ) appeared as compact rings and were centrally located as expected. In contrast, many of the cells expressing EcFtsZ^M225F^ were filamentous and contained multiple FtsZ rings or bands, which appeared uncompacted (**Figure 6E**). Next, we calculated the percentage of filamentous cells (defined as cells longer than 7 µm, (Haeusser *et al*., 2015)), of both wild type and M225F mutant in all the strains. The result showed higher percentage of cells (46.98 ± 2.84 for CCD288 at 37 °C, and 48.54 ± 18.93 for JKD7_1 strain expressing mutant EcFtsZ^M225F^ at 42 °C) with cell length above 7 µm. On the other hand, the wild type FtsZ expressing strains had a lower percentage of cells (i.e., 5.10 ± 3.92 for CCD288, and 10.56 ± 2.55 for JKD7_1 strain) longer than 7 µm (**Figure S4E**).

These results therefore suggest that the M225F mutation in EcFtsZ impairs the cell division of *E. coli* by affecting the Z-ring assembly and compaction of the ring.

## Discussion

There are two structurally different states of FtsZ that help in its cooperative assembly into single protofilaments - R (monomer) and T (filament) states (Caplan and Erickson, 2003; Matsui *et al*., 2012; Wagstaff *et al*., 2017). Based on *in vivo* observations, FtsZ is known to treadmill at the site of division wherein one end polymerizes and the other end depolymerizes (Bisson-Filho *et al*., 2017; Yang *et al*., 2017). The difference in the rate of assembly and disassembly between two ends of filaments, the basis for which is not completely understood, defines the kinetic polarity. The conformational switch from the R- to T-state is a defining feature which promotes preferential growth from one end, leading to the cytomotive behaviour of tubulin-like filaments (Wagstaff, et al, 2023). Our results support a structural basis for GTP-dependent polymerization preferentially to one end of the single protofilament and, hence, for the kinetic polarity exhibited by FtsZ filaments. The directionality of growth proposed by us is the same as that observed from the earlier study on FtsZ polarity (Du *et al*., 2018).

Based on the crystal structures of SmFtsZ, we propose that the Gly71 peptide bond of the T3 loop is re-oriented to accommodate the gamma phosphate, in the GTP- (or non-hydrolysable analog equivalents) bound structure. The loop is disordered when GDP is present, probably due to the missing interactions with the T3 loop. This is consistent with previous reports, which state that GTP binding stabilizes the T3 loop of FtsZ (Díaz *et al*., 2001; Yoshizawa *et al*., 2020; Ruiz *et al*., 2022). The flipping of the peptide favors the interaction with the next subunit as seen in the filament structure of FtsZ from *S. aureus* (**Figure 4F**;(Ruiz *et al*., 2022)) as well as the inferences from the DNP NMR study of GTP bound FtsZ filaments of *E. coli* (McCoy *et al*., 2022). Our structures represent conformations of the T3 loop captured in the same crystal packing environment with a disordered T3 conformation in the GDP-bound conformation and a preference for a flipped Gly71 peptide in the GMPPNP-bound state. The observation of a flipped peptide state of the T3 loop in response to GMPPNP-binding and our structural analysis from other FtsZ structures is suggestive of GMPPNP (or GTP) binding stabilizing the favorable conformation of T3 loop that assists in polymerization. We propose that the peptide flip and the stabilization of T3 loop could probably be deciding factors that determine the switch to a polymerization-competent monomer in the presence of GTP but not in GDP.

Our sequence and structure analysis showed that SmFtsZ has a phenylalanine in place of a flexible methionine on S7 beta strand in the interdomain cleft. We observe that the cleft mutant, SmFtsZ^F224M^ has higher GTPase activity than that of wild type. We infer that the mutation eases the transition of R- to T-state by facile opening of the cleft. The residue is located at the hinge where the inter-domain cleft opens. A flexible residue (Met224) in place of a rigid bulky residue (Phe224) might act as a lubricant for easier transition between states. The slower cleft opening in wild type SmFtsZ, therefore, could explain the higher CC for polymerization and lower GTPase activity of wild type SmFtsZ than that of EcFtsZ under identical *in vitro* conditions. In the *in vivo* study, when EcFtsZ-Met225 was mutated to phenylalanine, we observed filamentous morphology of cells indicative of aberrant cell division, possibly due to perturbation of FtsZ assembly leading to a cell division defect. It is likely that the slow R- to T-state transition due to the bulky side chain of phenylalanine in the cleft affects the rate of filament elongation. It is interesting to note that the mutant could form longer filaments though the GTPase activity was higher. Higher GTPase need not always be associated with shorter filaments, as observed in in vitro studies comparing *E.coli* and *Bacillus subtilis* FtsZ (Buske and Levin, 2012).

Finally, we propose a model for the kinetic polarity in FtsZ polymerization leading to the observed treadmilling behaviour. GTP binds R-state or the monomer state of FtsZ (evidenced by crystal structures where 25 FtsZ monomers have been captured with GTP or analogs in the R-state conformation) (**Figure 7A, Figure S5 step 1**). The GTP-bound NTD presents a pre-formed conformation for binding at the bottom end of the filament (Ruiz, et al, 2022). In addition to the Mg^2+^ ion coordination and the active site stabilization with the gamma phosphate (Ruiz, et al, 2022), the flipped out Gly71 peptide could establish inter monomer contacts within FtsZ filament (**Figure 4F**). Attachment of a monomer to the bottom of the filament requires the downward movement of its H7 helix (**Figure 7B, case (ii); Figure S5 step 2**), which is a characteristic of the T-state. The downward movement of the H7 helix is accompanied by the repositioning of Phe183 and rotation of the guanine ring (**Figure 7B, case (ii); Figure S5 step 3**). Downward position of the H7 helix in the T-state clashes with the beta sheet of the CTD in the R-state (**Figure 7B, case (ii); Figure S5, step 3**). Hence, this might trigger the opening of the cleft, resulting in a complete conformational change and stabilization of the T-state (**Figure 7B, case (ii); Figure S5, step 4**).

**Figure 7.**
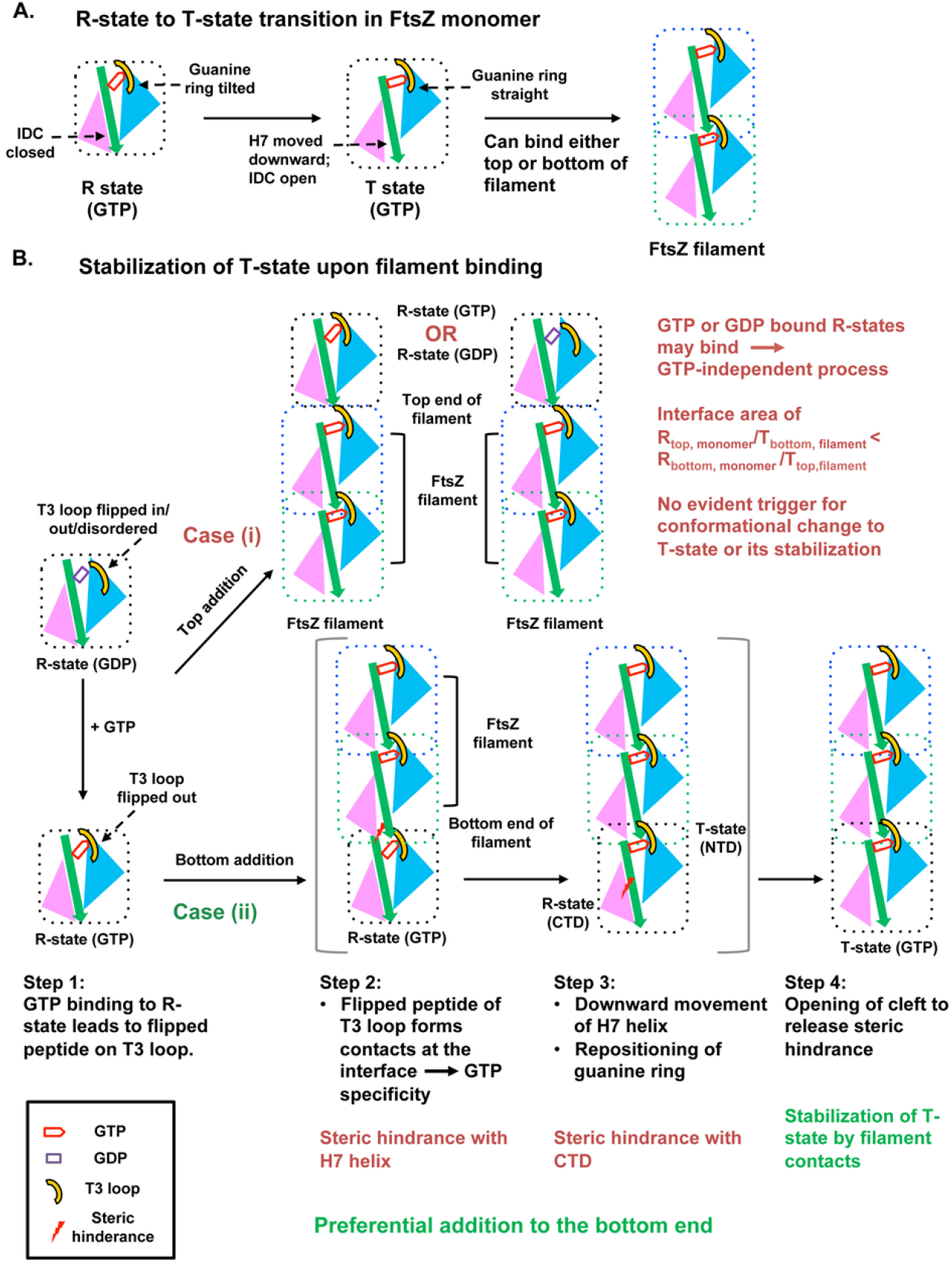
Model for FtsZ polymerization and kinetic polarity. **A. R- to T-state transition in FtsZ monomer.** If a T-state FtsZ monomer exists in solution, it could bind to either top or bottom of the filament without any preference to one of the ends. In such a scenario, there will be no structural or kinetic polarity in the process of polymerization of FtsZ filament. **B.** Stabilization of T-state upon filament binding. **Case (i). Addition of R-state FtsZ (GDP/GTP) to top end of filament**: The R-state FtsZ irrespective of the identity of the bound nucleotide could bind to the top end of the filament via the closed interdomain cleft (IDC). There is no trigger for the downward movement of H7 helix as the H7 helix in R-state is upwards and the addition is irrespective of nucleotide. Therefore the top addition for filament growth might be less feasible. **Case (ii). Addition of R-state FtsZ (GTP) to bottom end of filament** **Step 1**: **GTP binding to R-state and flipping out of T3 loop**: The monomer state (R-state) of FtsZ binds GTP. The GTP binding leads to flipping out of the Gly71 peptide and also the gamma phosphate directly interacts with the T3 loop. **Step 2**: **Binding to bottom subunit via flipped out T3 loop**: The GTP bound R-state FtsZ can, in principle, attach to the bottom interface of the FtsZ filament via the T3 loop. **Step 3**: **Downward movement of H7 helix, repositioning of guanine ring and clash with CTD of closed IDC**: In order to accommodate the FtsZ monomer at the bottom of filament, H7 helix has to move downwards. The downward movement of the H7 helix repositions Phe183, which forms stacking interaction with the guanine ring of the nucleotide. This leads to the repositioning of the guanine ring, which is observed in T-state structures. As the H7 helix moves downwards, there are clashes between the residues of the H7 helix and the beta sheets of CTD in the closed IDC (interdomain cleft). **Step 4**: **Cleft opening and conversion to T-state**: The clashes in the interdomain cleft trigger the opening of the cleft and changing the entire molecule to T-state. The conversion of T-state occurs upon binding to the filament, only after GTP binding. The NTD is in sky blue; the H7 helix and the T7 loop is denoted in green; the CTD is in pink; the purple rectangle denotes GDP and the red pentagon denotes GTP.

The efficiency of the opening of the cleft depends on the ease of rearrangement of interactions between the central H7 helix and beta strands S8, S9, S10 including the S7 beta sheet. A flexible residue in S7 beta strand at the interdomain cleft is important for the smoother transition from R- to T-state favoring more efficient filament formation and hydrolysis. If the T-state conformation were stable for FtsZ monomers in solution, cooperativity for addition to the single protofilament or kinetic polarity might not be very pronounced, as a pre-formed T-state monomer can, in principle, bind to both the ends with equal chance (**Figure 7A**). In summary, our model provides an explanation of how GTP binding can trigger a conformational change to the T-state upon binding to the bottom end of the filament. However, it appears that the transition to the T-state may not be as efficient upon addition to the top of the filament. The addition of a monomer to the top end of the filament might be less favoured than the addition at the bottom of FtsZ filament. The buried surface area between the closed cleft of incoming subunit and the NTD of the filament (interacting interfaces for top addition) is lower than the buried surface area between the opened cleft of the filament and the NTD of the incoming subunit bound to GTP (interacting interfaces for bottom addition) (Lv *et al*., 2021). When added on to the top, the C-terminal domain end should drive the conformational change to the T-state to improve interprotomer contacts, which might be a rate limiting step given that the closed interdomain cleft can be accommodated without any steric clashes. Additionally, the discriminatory role of GTP becomes irrelevant upon addition of a monomer from the top, which might occur irrespective of the nucleotide state (**Figure 7B, case (i)**), especially relevant in the context of single protofilament growth as in FtsZ. Lateral interactions might play a role in this discrimination in a multi-protofilament scenario like in microtubules.

The crystal structures of wild type as well as the cleft mutant of SmFtsZ show domain swap dimers. It is possible that there is an inherent tendency for the NTD and CTD to go into an extended conformation if addition of subsequent monomers does not stabilize the T-state conformation. The extended conformation is not stable and thereby interacts with another monomer in extended conformation to form a domain-swapped dimer, as observed in crystal structures of wild type SmFtsZ and non-polymerizing mutants of other FtsZs too (PDB IDs: 1W5F, 5MN7; (24, 45)). In our crystal structures, the polymerization-incompatible domain-swapped dimers of SmFtsZ might preferentially crystallize compared to the globular monomers which tend to polymerize. The presence of strong contacts by means of a phenylalanine residue at the inter-domain cleft of SmFtsZ could act as a barrier for a smooth transition to the T-state. SmFtsZ monomer might have a higher propensity compared to EcFtsZ to go through an extended conformation during conversion from R-state to T-state. Previous reports have indeed shown that the NTD and the CTD are independent of each other, which can reconstitute non-covalently to form functional FtsZ (Oliva *et al*., 2004), similar to a domain-swapped globular arrangement of NTD and CTD.

Our observations highlight the parallels in the polymerization mechanism of tubulin and FtsZ. In contrast to FtsZ, tubulin is composed of a heterodimer, which polymerizes to form microtubules (Nogales *et al*., 1998). It is interesting to speculate that the existence of tubulin as a heterodimer has probably evolved to prevent a non-productive extended conformation of the NTD and CTD of the tubulin fold.

## Materials and Methods

### Cloning

*ftsZ* genes from *S. melliferum* (UniProt ID: A0A4D8RL11) and *E. coli* (Uniprot ID: P0A9A6) were amplified from the genomic DNA of the respective organisms (*S. melliferum*: genomic DNA obtained from DSMZ, Germany, catalogue number 21833; *E. coli*: genomic DNA extraction from lab strain). In *Spiroplasmas*, the codon TGA codes for tryptophan whereas in *E. coli*, TGA is a stop codon. Therefore, for using *E. coli* as the heterologous expression system, all TGA codons in the *ftsZ* gene were changed to TGG (the codon which codes for tryptophan in *E. coli*). *ftsZ* gene from *S. melliferum* has two tryptophans which are coded by TGA, at the amino acid positions 384 and 411. These two tryptophan codons were changed to *E. coli* codon (TGG) using primers for site-directed mutagenesis. The genes were cloned into pHis17 vector (obtained from Löwe lab, MRC LMB, Cambridge; refer Addgene plasmid #78201 for vector backbone) using restriction-free cloning (van den Ent and Löwe, 2006) between *NdeI* and *BamHI* restriction enzyme sites. The truncated constructs and the single-point mutants were also generated using PCR followed by *DpnI* digestion, transformation and screening for the positive clones using sequencing. pCCD1020, carrying the EcFtsZ mutant M225F under the arabinose promoter was created using restriction-free cloning in pCCD436 (pJSB100; pBAD18Cm-EcFtsZ) (Stricker and Erickson, 2003) by using primers RSO1107 and RSO428. pCCD1042 was constructed by PCR amplification of SmFtsZ using primers RSO1081 and RSO1083 and replacing EcFtsZ from pCCD436 by restriction-free cloning. pCCD1043 (SmFtsZ_1-314_-mNG-MinD_MTS_) carries the core 1 – 314 amino acid residues of SmFtsZ fused to mNeonGreen with the EcMinD_MTS_ at its C-terminus. The construct was created by restriction free cloning after amplification of SmFtsZ_1-314_ by PCR using primers RSO1081 and RSO1082, replacing EcFtsZ_1-366_-mNG-MinD_MTS_ in pJSB2 plasmid (pCCD907). pCCD907 is identical to the EcFtsZ_1-366_-Venus-MinD_MTS_ reported by Erickson and colleagues (Osawa *et al*., 2009) and its construction will be reported elsewhere. The list of primers used for cloning of the above genes and list of strains and plasmids used in this study are given in **Tables S6, S7, S8**.

### Protein purification

The plasmids were transformed into *E. coli* BL21(AI) cells and 1 L culture was grown in LB medium supplemented with ampicillin. The cultures were induced with 0.2 % (w/v) L-arabinose when its optical density (OD) at 600 nm reached 0.6 and was further incubated at 30 °C for 5 hours post-induction and cell harvested by spinning at 6000 *xg* at 4 °C. The pellet was resuspended with lysis buffer (50 mM Tris, pH 8, 200 mM NaCl and 10% glycerol) and sonicated. The lysate was centrifuged at 100,000 *xg* at 4 °C and the supernatant was loaded onto a 5 mL His-Trap (GE Lifesciences) column pre-equilibrated with buffer A (50 mM Tris, pH 8, 200 mM NaCl). The protein was eluted with a step gradient of imidazole containing buffer B (50 mM Tris, pH 8, 200 mM NaCl, 500 mM imidazole). The purest fractions of the protein were identified using 12 % SDS-PAGE gel, pooled, concentrated and injected into the gel filtration column, Superdex 200 10/300 (GE Lifesciences), pre-equilibrated with the final buffer (HK50-50 mM HEPES, pH 6.5, 50 mM KCl). The protein eluted at volumes corresponding to monomeric (major peak) and dimeric (small hump) size for SmFtsZ constructs. The fractions were pooled, concentrated, flash-frozen in aliquots, and stored at -80 °C till further use. The same protocol was followed for all other constructs of FtsZs. For wild type EcFtsZ, only affinity chromatography was carried out. The final proteins were visualized in SDS-PAGE gel to check for the purity (**Figure S1A**). The molecular weight of the purified wild type SmFtsZ was confirmed using mass spectrometry.

### Pelleting assay for estimating filament content

Protein was spun at 100,000 *xg* at 4 °C in order to remove any precipitate. 10 μM of the protein was taken from the supernatant to set up 50 μL of reaction. The following reaction conditions were prepared-a) only protein, b) protein and MgCl_2_, c) protein, MgCl_2_ and GDP, d) protein, MgCl_2_ and GTP, e) protein, MgCl_2_ and GMPPNP (GNP), and protein, MgCl_2_ and GTPγS (GSP). A final concentration of 3 mM of nucleotide and 5 mM of MgCl_2_ was taken for the reactions. The reaction mixtures were incubated at 30 °C for 10 minutes, spun at 100,000 *xg* at 30 °C for 15 minutes. The supernatant was taken out and the pellet was resuspended with a buffer as equal volume of the reaction. The supernatant and pellet fractions were mixed with 2x SDS buffer and loaded onto 12 % SDS-PAGE gel in equal amounts.

### Malachite green assay for estimating released phosphate to measure GTPase activity

The GTPase activity of all constructs was performed using Malachite green assay which is a colorimetric assay and measures the release of inorganic phosphate (Feng *et al*., 2011). The protein was spun at 22,000 *xg* at 4 °C and the reaction was set with the required concentration of protein, reaction buffer (50 mM HEPES, pH 6.5, 50 mM KCl), saturating concentration of GTP (obtained from Michaelis-Menten plot, EcFtsZ-1mM, SmFtsZ-2mM), and 5 mM MgCl_2_.

The reaction mixture (20 μL) was incubated at 30°C for 15 minutes and stopped by addition of 100mM of EDTA (5 μL of 0.5M of EDTA). The reaction mixture was added to a 96 well plate and 100 μL of the malachite green mixture was added into the reaction mixture. Also, a set of phosphate standard solutions of known concentrations was prepared and added to the 96 well plate along with the malachite green mixture. After 10 minutes of incubation, absorbance was measured in Varioskan Flash (4.00.53) at 630 nm wavelength.

The concentration of phosphate released was calculated by dividing the slope obtained from the phosphate standard. The reaction containing buffer and only GTP was taken as blank and the absorbances were normalized by subtracting the blank. The Michaelis-Menten plots were generated with the y-axis as the rate of reaction that is phosphate released per unit time (μM/min) and the x-axis as different concentrations of GTP from 0-2.5 mM for EcFtsZ and 0-10 mM for SmFtsZ. The curve was fitted with the Michaelis Menten equation, V_0_ = V_max_ ([S]/([S] + K_M_).

The *k_obs_* (min^-1^) of the proteins were calculated by dividing the concentration of phosphate released by time and the concentration of the protein and *k_cat_* (min^-1^) was determined by the formula V_max_/[E_t_] from the Michaelis-Menten plot. The critical concentrations of the proteins were determined by obtaining the *k_obs_* (min^-1^) of different concentrations of the protein ranging from 0 to 10 μM for EcFtsZ and 0 to 14 μM for SmFtsZ. Statistical analysis was done with one-way ANOVA test and the statistical significance values were obtained using Tukey’s multiple comparisons test. All the statistical analysis and plotting was done using version 9.00 for MacOS of GraphPad Software (San Diego California USA). Mean and standard error of mean (SEM) were calculated.

The value of the specific activity of EcFtsZ was comparable with the previously reported values (Mukherjee, 1998; Dajkovic *et al*., 2008; Mohammadi *et al*., 2009), but higher values were also reported by other studies (Dai and Lutkenhaus, 1991; de Boer *et al*., 1992; Mukherjee *et al*., 1993; Wang and Lutkenhaus, 1996; Yang *et al*., 2017). The differences in the values of the specific activity of EcFtsZ might be due to differences in purification protocol, method for determining phosphate release, reaction buffer, and temperature.

### Size exclusion chromatography (SEC) and Size exclusion chromatography with multi-angle light scattering (SEC-MALS)

For SEC-MALS, 100 μL of 2 mg/mL SmFtsZ^ΔCT^ was injected in GE superdex 200^TM^ column equilibrated with HK50 buffer (50 mM HEPES, pH 6.5, 50 mM KCl). The apparatus was connected to Wyatt Heleos II 18° angle light scattering which is coupled to a Wyatt Optilab rEX online refractive index detector. The data analysis was done using the ASTRA software 6.1.7.17 and the molar mass of the injected protein was calculated from the differential refractive index and the light scatter.

For SEC, 200 μL of 10 mg/mL SmFtsZ^ΔCT^ was injected in GE superdex 200^TM^ column equilibrated with HK50 buffer. The data was plotted in 9.00 for MacOS, GraphPad Software.

### HPLC run to determine the nucleotide bound to the protein

Purified SmFtsZ was diluted to 176 μM in HK buffer (pH-6.5) and was denatured at 95 °C for 1 minute, spun at 21,000 *xg* for 10 mins at 4 °C. The supernatant was filtered using 0.22 μm cellulose acetate filter (Corning) and the sample was loaded on a DNAPac PA 200 ion exchange analytical column pre-equilibrated by buffer A (2 mM Tris, pH 8). The runs were performed using a linear gradient of 0 – 50 % buffer B (2 mM Tris, pH 8, and 1.25 M NaCl) for 5 column volumes (CV), 50 – 100 % buffer B for 3 CV and 0 % buffer B for 1 CV. The runs were performed at 0.5 ml/min. The peak profile obtained was compared for runs of 7 nmoles and 10.6 nmoles of SmFtsZ (stock concentration-176 uM). The standards used for the run were GDP of 6 nmoles, 12 nmoles, 24 nmoles, 36 nmoles, 48 nmoles, and 60 nmoles from a stock of 100 mM. The graph was plotted with absorbance at 254 nm against the retention time using version 5.00 for Windows, GraphPad Software.

### Crystallization of SmFtsZ^ΔCT^ with and without nucleotide addition

SmFtsZ^ΔCT^ crystallization was set up using around 500 commercially available screens (Molecular dimensions, Hampton Research) having 100 nL of 10 mg/mL protein and 100 nL of the respective condition in 96 well sitting drop plates where drop A had only protein with the condition and drop B had protein, GMPPNP, MgCl_2_, and conditions. The initial hits obtained were optimized in order to get well diffracting crystals. Needle-like crystals were obtained in the conditions tabulated in **Table S9**. The crystals were frozen with 20 % (v/v) ethylene glycol as cryoprotectant in the parent condition.

### Structure determination

The crystals were diffracted and the final data collected at synchrotron sources at Diamond Light Source (Harwell, UK) and ELETTRA (Trieste, Italy). The diffraction data was indexed using XDS and scaled using AIMLESS present in the CCP4i package (Evans and Murshudov, 2013; Potterton *et al*., 2018). Molecular replacement was done with BsFtsZ structure (PDB ID: 2VXY) using PHASER and model building was done using COOT (McCoy *et al*., 2007; Emsley *et al*., 2010). Refinement was done using PHENIX (Adams *et al*., 2010). The data collection and refinement statistics are tabulated in **Table S3**.

### Transmission electron microscopy

Transmission electron microscopy was carried out to visualize the filaments of the wild-type SmFtsZ and SmFtsZ^F224M^. The protein was spun at 100,000 *xg* at 4 °C and a reaction of 20 μL was set with a final protein concentration of 10 μM and GTP concentration of 3 mM at 30 °C for 15 minutes. Another set of samples was prepared by resuspending the pellet obtained after 100,000 *xg* spin for 15 minutes at 30 °C of the aforementioned reaction, with the same volume of buffer as the reaction.

5 μL of protein was added on a glow discharged (one round of 15 mA current for 25 seconds using plasma cleaner, Quorum Technologies) carbon coated copper grid and incubated for 2 minutes before blotting out the protein. 5 μL of 1.5 % (w/v) uranyl acetate was added and incubated for 30 seconds and blotted out. The grids were dried before imaging under TEM (JEM-2200FS Joel Ltd.). The images of the filaments were analysed using ImageJ (Schneider *et al*., 2012). 2-D class averages were obtained using default options in CryoSPARC software (Punjani *et al*., 2017).

### Sequence analysis

FtsZ sequence of *E. coli* (UniProt ID: P0A9A6) was used to acquire sequences using the EVfold webserver (Hopf *et al*., 2019). The sequences with identity percentages ranging from 19 – 100 % were retrieved from UniProtKB. All significant hits except redundancies and low-quality sequences were selected and aligned with MUSCLE in JALVIEW (ver 2.11.4) (Waterhouse *et al*., 2009). After generating the alignment, a 90 % redundancy cutoff was applied. The total number of sequences considered for the analysis was 818.

For acquiring only *Spiroplasma* FtsZ sequences, the FtsZ sequence from *S. melliferum* was used as a query sequence for BLAST against all *Spiroplasmas*. A total of 26 *Spiroplasma* FtsZ sequences were obtained.

### Structural analysis

For the analysis, all the available structures of FtsZ structures are used, as given in **Table S4**. For all the superimpositions, the NTD of the molecules (residue number 1 to 170) was superimposed using the ‘matchmaker’ command in Chimera. All the analyses are done using UCSF Chimera version 1.13.1 (Pettersen *et al*., 2004).

### Growth and culturing of bacteria

*E. coli* strain (CCD288) (*BW27783 FtsZ 55-56 SW-mNG*) was grown in LB Lennox at 37 °C. CCD56, *E. coli* strain JKD7_1/ pKD3 (*repA^ts^*) (35) was grown in LB Lennox containing 25 μg/ mL Kanamycin and 100 μg/ mL carbenicillin at 30 °C or at a restrictive temperature of 42 °C to allow for the loss of pKD3 plasmid. Chloramphenicol, wherever necessary, was used at a concentration of 34 μg/ mL. Expression of EcFtsZ, EcFtsZ^M224F^, SmFtsZ from pCCD434 (pJSB2) derived plasmids was achieved by the addition of 0.2 % L-arabinose. Glucose at 1 % was used for repression. *E. coli* strain, CCD161 (BW27783) was transformed with plasmids pCCD907 (EcFtsZ_1-366_-mNG-MinD_MTS_) or pCCD1043 (SmFtsZ_1-314_-mNG-MinD_MTS_) and grown at 37 °C. Protein expression was achieved by the addition of 0.1 % arabinose and growing the cultures further for 2 hours at 37 °C before imaging. Where mentioned, cephalexin (10 µg/ml) was used to inhibit cell division and obtain filamentous cells for imaging.

### Complementation and Spot Assays

Complementation assay was performed as previously described by (Stricker and Erickson, 2003). In the strain *E. coli* JKD7-1 carrying the plasmid pKD3, the chromosomal *ftsZ* gene is disrupted by a kanamycin cassette. EcFtsZ is expressed from the plasmid pKD3, which is temperature sensitive for replication due to the *repAts* allele. Thus, the growth of this strain is permissible only at 30 °C as the plasmid ceases to replicate at higher temperatures, resulting in the loss of *ftsZ*. However, at 42 °C, growth can be achieved only upon supplementing FtsZ from an additional plasmid such as pJSB100, wherein FtsZ is expressed from an arabinose inducible promoter.

Briefly JKD7-1/pKD3 strain carrying pCCD434 or pCCD434 derived plasmids (pCCD436, pCCD1020 and pCCD1042) was grown overnight in repression medium containing 1 % glucose at 30 °C. The overnight culture is serially diluted starting with OD_600_ 0.8 - 1.0 and then increasing dilution factor from 5^0^ to 5^6^. Three microliters of each dilution were spotted on repression plates containing 1 % glucose and induction plates containing 0.2 % L-arabinose using a 48-pin replicator (VP-Scientific). The plates containing 0.2 % L-arabinose were incubated at a restrictive temperature of 42 °C and repression plates containing 1 % glucose at 30 °C. *E. coli* strain (CCD288; *BW27783 FtsZ 55-56 SW-mNG*) carrying pCCD434 derived plasmids (pCCD436, pCCD1020 and pCCD1042) was grown at 37 °C and similarly spotted in LB plates containing either 1 % glucose or 0.2 % L-arabinose and incubated for 14 – 16 hours at 37 °C.

### Western Blot

Expression of FtsZ was detected by western blotting using anti-FtsZ antibodies (AS10715: Agrisera). CCD288 was grown as described above and cell lysates were subjected to SDS-PAGE, followed by western blotting. Briefly, 3 mL cultures were harvested by centrifugation at 2800 *xg* for 5 minutes, followed by 3 times wash with 1X PBS and resuspended in a 100-200 μL Laemmli buffer with 10-20 μL β-mercaptoethanol. The suspension was vortexed and left on ice for 5 minutes. The samples were then sonicated (40 % amplitude with 1 second on & 3 seconds off) for 3 - 4 cycles. The lysate was centrifuged at 10,000 *xg* for 2 minutes and supernatant was transferred into a fresh tube and 1.8 μL of Laemmli buffer was added and heated at 98 °C for 10 - 15 minutes, before loading the samples to 12 % SDS-PAGE gel. The proteins in the gel were transferred to a polyvinylidene difluoride membrane (Immobilon-P) using a semidry transfer apparatus (Amersham Life Sciences or Trans Blot Turbo Transfer System from Bio-Rad). Membranes were blocked using PBS containing 2% BSA and 0.05% Tween-20. Subsequently, anti-FtsZ antibody (AS10715: Agrisera) at a dilution of 1:10,000 was prepared in 1X PBS containing 2% BSA and 0.05% Tween-20 and incubated for 1.5 h at room temperature (RT). The blots were washed three times in PBST (0.05% Tween-20). Then Protein A-HRP at a dilution of 1:15,000 prepared in 1X PBS containing 2% BSA and 0.05% Tween-20 was added and incubated for 1 hour at RT. The blot was subsequently washed 3-4 times with 20 mL of PBST (0.05% Tween-20). Final detection was carried out using chemiluminescence HRP substrates (Immobilon Western Chemiluminescent HRP Substrate; WPKLS0500; Merck Millipore) and imaged using ChemiDoc XRS+ (Bio-Rad).

### Microscopy

The bacterial strains (JKD7-1/pKD3 with appropriate plasmids) were grown overnight in repression media at 30 °C and then sub-cultured by diluting 1:100 times into media containing 0.2 % L-arabinose. Growth was continued further at 42 °C by diluting 1:10 times at 1.5 hours intervals at least twice. The cultures were grown to an OD_600_ between 0.3 - 0.6- and 1.5-mL culture was centrifuged at 2800 x *g* for 3 minutes. Microscopy was performed similar to what has been shown in (Sharma *et al*., 2023). The cell pellets were gently resuspended in 150 µL of fresh LB medium of which 1 – 2 µL were spotted on 1.7 % LB-agarose pads and imaged. Similarly, CCD288 strain was grown overnight at 37 °C, diluted 1:100 into fresh medium and imaged using agarose pads after 2 hours when they reached an OD_600_ of 0.4 – 0.6. The cells were imaged on agarose pads. LB-agarose pads were prepared with LB (Lennox) media containing 1.7 % agarose in cavity slides and covered with a coverslip No. 1.5 (VWRI631-0125). Images were acquired using an epifluorescence microscope (DeltaVision^TM^ Elite) with a 100X oil immersion phase objective (PLN100XOPH) of NA 1.25 and an in-line CCD camera (CoolSNAP^TM^ HQ2). An oil of refractive index 1.514 was used for imaging bacteria cells. A Spectra 7, Lumencor Light Engine^©^ (solid-state illumination) was used as a light source with a 1 mm diameter liquid light guide. Excitation filters and emission filters of 488/28 nm and 525/48 nm respectively were used for imaging mNeonGreen tagged strains.

### Three dimensional-Structured Illumination Microscopy (3D-SIM)

Cells were mounted on an agarose pad as described above and imaged using a DeltaVision^TM^ OMX-SR Blaze microscope. To better visualize the polymers assembled by FtsZ tagged with mNeonGreen tethered to the membrane, filamentous cells were obtained by treating the cultures with 10 µg/ml cephalexin for 2 hours to inhibit cell division. For 3D-SIM, 15 images (5 translations x 3 rotations) were obtained for each z-plane. The section spacing of 0.125 µm to cover the width of the bacterial cell (*E. coli*) around 2 µm was used for Z-sectioning. Images were acquired using a 100x, NA 1.42 oil-immersion objective lens and a PCO Edge 4.2 sCMOS camera. An exposure time of around 10 – 15 ms and % transmittance of 20% was used to acquire each image. 3D-SIM were reconstructed using the SI reconstruction module of SoftWorx^TM^ software using deconvolution, quick projection and SIM reconstitution options. For imaging the mNeonGreen-tagged strains, excitation and emission filters of 488/28 nm and 525/48 nm respectively were used.

## Author Contributions

JC conceptualized, designed and performed all the experiments, analyzed data other than those mentioned below and wrote the manuscript, SMP performed, analyzed and wrote the *in vivo* section, SD helped in cloning and purification of the SmFtsZ^D211G^ mutant, 2D class average analysis and edited the manuscript, VB performed the sequence analysis, assisted with protein purification, pelleting assay and HPLC, SH assisted in standardization of purification and pelleting assays of EcFtsZ, RS conceptualized, supervised and wrote the *in vivo* work, PG initialized, conceptualized, supervised and managed the project and wrote the manuscript.

## Classification

Biochemistry, Microbiology

## Acknowledgments

Synchrotron facilities at Diamond Light Source, UK and ESRF, Grenoble [through ESRF Access Program of the Department of Biotechnology (Grant No. BT/INF/22/SP22660/2017)] and access to DST ELETTRA XRD2, Jyotsna from Dr. Radha Chauhan’s lab at NCCS for access to the SEC-MALS facility at NCCS, Pune, macromolecular crystallography facility at IISER Pune, and electron microscopy facilities at the Advanced Technology Platform Centre, Regional Centre for Biotechnology, Faridabad, and at IISER Pune with support from DST under Nanomission program (EMR/2016/003553) are acknowledged. We acknowledge Prof Jayant Udgaokar’s lab in IISER Pune for help in mass spectrometry. Centre of Interdisciplinary Sciences, NISER, for OMX Blaze-SR and its support staff member Mr. Mirza Salim Beg are acknowledged. We thank Mr. Ajay Kumar Sharma for the construction of the plasmid pCCD907. We thank Dr. Harold Erickson (Duke University, USA) and Dr. Kerwyn Casey Huang (Stanford University) for strains and plasmids. We acknowledge fellowships from IISER Pune to JC and SH, DAE to SMP, and CSIR to SD. Research work in the lab of PG was supported by Department of Biotechnology (DBT) Membrane Structural Biology Program grant (BT/PR28833/BRB/10/1705/2018), SERB CRG (CRG/2018/003795) and IISER Pune, and SERB (CRG/2021/000337), DBT (BT/INF/22/SP33046/2019) and intramural core funding from Department of Atomic Energy (DAE) to RS.

