## Supplemental files for "Dynamics of interdomain rotation facilitates FtsZ filament assembly"

**This PDF file includes:**

Figures S1 to S5
Tables S1 to S9
Reference

Figure S1

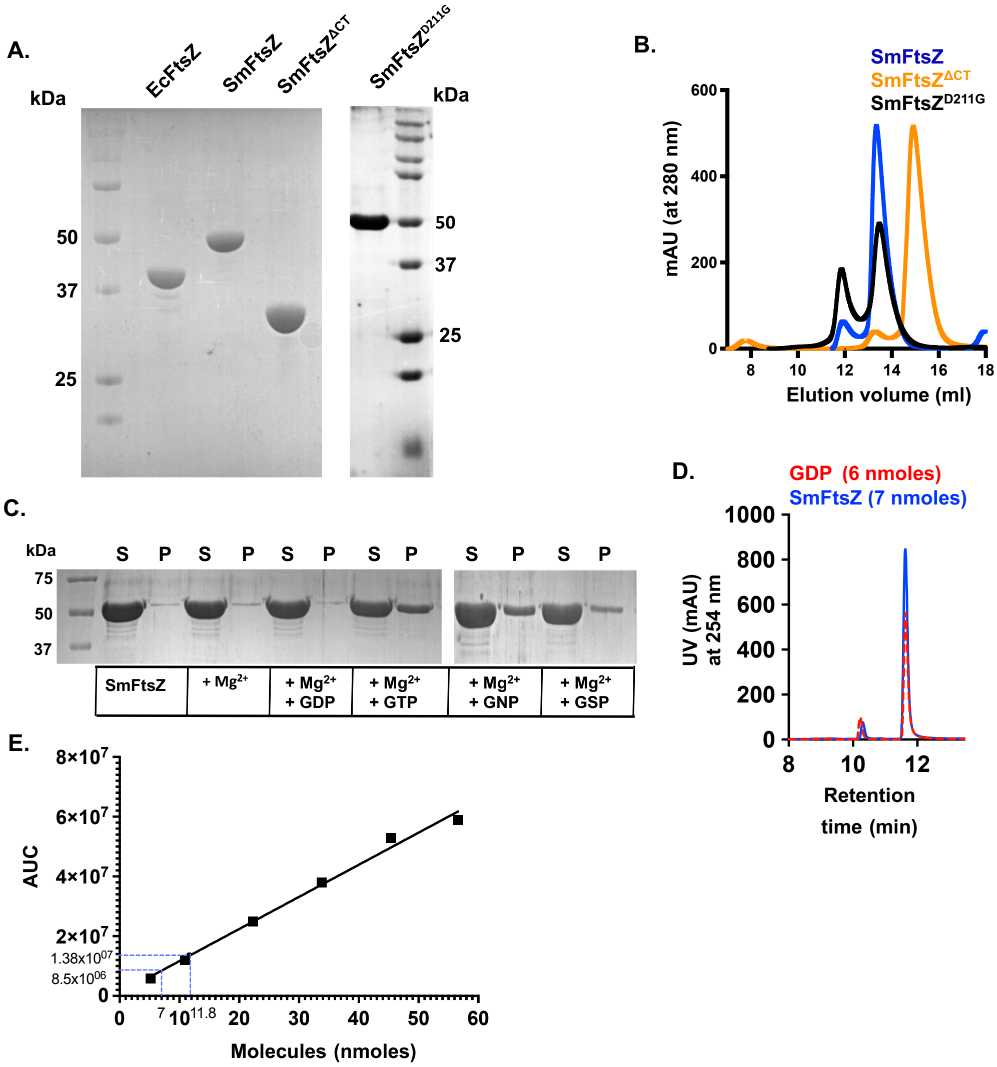


**Figure S1. Biochemical characterization of SmFtsZ**

**A)** A representative 12% SDS-PAGE gel image denoting the purified proteins used in the study- EcFtsZ, SmFtsZ, SmFtsZ$^{\Delta CT}$ and SmFtsZ$^{D211G}$. **B)** Analytical run in Superdex200 column showing the oligomeric status of the proteins. For SmFtsZ and SmFtsZ$^{D211G}$, the elution volume is 12 ml (minor peak- dimer) and 13.4 ml (major peak- monomer). For SmFtsZ$^{\Delta CT}$, the elution volume is 13.5 ml (minor peak- dimer) and 15 ml (major peak- monomer). **C)** A representative SDS PAGE gel for pelleting assay of SmFtsZ to determine the condition necessary for SmFtsZ filament formation in pH 6.5. Incubation of SmFtsZ (10 μM) in presence of $Mg^{2+}$ (5 mM); $Mg^{2+}$ (5 mM) and GDP (3 Mm); $Mg^{2+}$ (5 mM) and GTP (3 mM); $Mg^{2+}$ (5 mM) and GMPPNP (GNP) (3 mM); and $Mg^{2+}$ (5 mM) and GTPγS (GSP) (3 mM). The presence of SmFtsZ in the pellet suggested that it formed filaments. S and P represent the fraction in supernatant and pellet of the reaction. (*N* = 2, *n* = 6). **D)** Plot for comparing the pellet fractions of the pelleting assay. The y-axis shows the percentage of the pellet fraction. (*N* = 2, *n* = 6). The error bar shows mean with SEM. **E)** HPLC run in DNAPac PA 200 column for 7 nmoles (stock concentration 176µM) of denatured SmFtsZ supernatant fraction (blue solid line) shows absorbance at 255 nm (milli absorbance unit [mAU]) of bound GDP peak (red dotted line) compared to standard GDP run (6nmoles). **F)** HPLC runs in DNAPac PA 200 of GDP standards for 6 nmoles, 12 nmoles, 24 nmoles, 36 nmoles, 48 nmoles, 60 nmoles from a stock concentration of 100mM. Two repeats of SmFtsZ 7 nmoles and 10.6nmoles (dotted line) extrapolated from the standard curve (Right).

### Figure S2

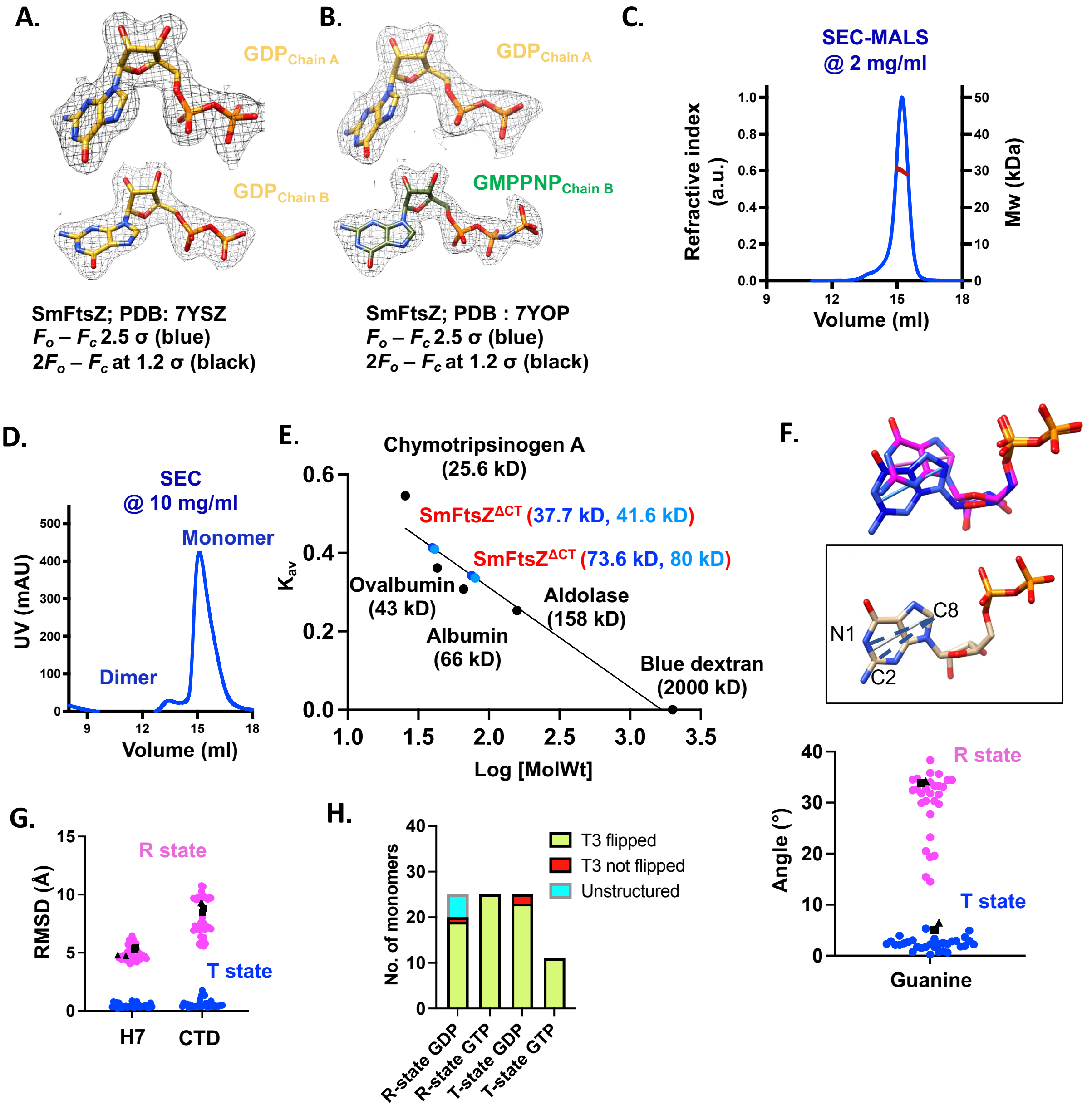


**Figure S2. Structure and oligomeric status of SmFtsZ**

**A)** Electron density map for the nucleotide in both the chains of SmFtsZ (PDB ID: 7YSZ). **B)** Electron density map for the nucleotide in both the chains of PDB ID: 7YOP. Map $F_o - F_c$ shown at 2.5 σ (blue) and $2F_o - F_c$ at 1.2 σ (black). **C)** SEC-MALS chromatogram of SmFtsZ $^{\Delta CT}$. SmFtsZ$^{\Delta CT}$ loaded at 100 μL of 2 mg/mL (low concentration) (protein concentration on the column is around > 1/10$^{th}$ of the concentration loaded). The profile shows a single monodisperse peak with average molecular mass at 30 kD. **D)** SEC profile of SmFtsZ$^{\Delta CT}$ at 200 μL of 10 mg/mL (high concentration) shows two peaks: one major peak at 15 mL (monomer) and another small hump at 13.5 mL (dimer). **E)** The calibration curve for size-exclusion chromatography for Superdex 200 using molecular weight standards shows that SmFtsZ elutes as monomer (major peak) and dimer (hump). The theoretical and estimated molecular weights of SmFtsZ $^{\Delta CT}$ are mentioned (Blue and light blue denote the repeats). **F)** Quantification of the guanine ring tilt angle keeping a T state structure (PDB ID- 5H5G) as reference. Inset shows atoms selected (C2, C8, N1) for guanine/nucleotide angle calculation. The pink cluster shows the structures in R state and the blue cluster shows the structures in T state. **G)** Quantification of the CTD movement, keeping T state structure (PDB ID-5H5G) as reference. The pink cluster shows the structures in R state and the blue cluster shows the structures in T state. Black triangle represents GMPPNP bound SmFtsZ structure (PDB ID: 7YOP) and black square represents GDP bound SmFtsZ (PDB ID: 7YSZ). **H)** Plot of FtsZ monomers in R- state GDP, R- state GTP, T- state GDP and T- state GTP and the orientation of Gly71 (numbering corresponding to SmFtsZ) in the T3 loop. Lime color shows flipped out conformation, the red color shows monomers where the peptide is not flipped out and the cyan color shows monomers where the T3 loop is unstructured.

### Figure S3

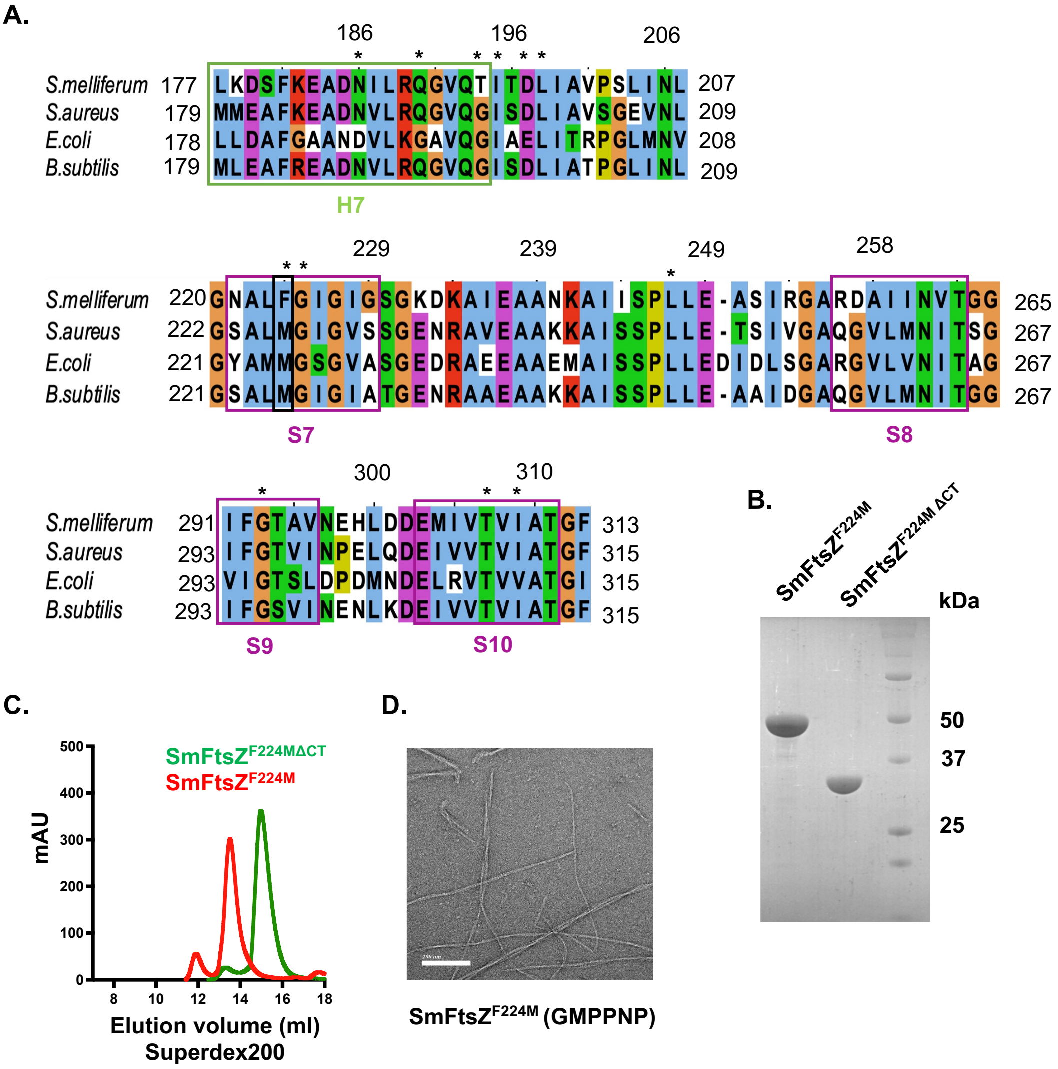


**Figure S3. Mutation in the inter-domain cleft of SmFtsZ.**

**A)** Multiple sequence alignment showing the residues involved in interdomain cleft interaction. The FtsZ sequences of *S. melliferum*, *S. aureus, E. coli*, and *B. subtilis* are shown as representative. Numbering of the residues is according to the FtsZ sequence from *S. melliferum*. The color code is according to the convention followed in ClustalX: Blue - hydrophobic, red - positive charge, magenta - negative charge, green - polar, pink - cysteine, orange - glycine, yellow – proline, cyan - aromatic. **B)** The 12% SDS-PAGE gel showing the final proteins of the interdomain cleft mutant- SmFtsZ$^{F224M\Delta CT}$ and SmFtsZ$^{F224M}$. **C)** Analytical run in Superdex200 column showing the oligomeric status of the proteins. For SmFtsZ$^{F224M}$, the elution volume is 12 ml (minor peak- dimer) and 13.6 ml (major peak- monomer). For SmFtsZ$^{F224M\Delta CT}$, the elution volume is 13.4 ml (minor peak- dimer) and 15.4 ml (major peak- monomer). **D)** TEM image of SmFtsZ$^{F224M}$ with 10 μM of protein and 2 mM GMPPNP. The experiments were done with saturating concentration of GTP. Scale bar represents 200 nm.

**Figure S4**

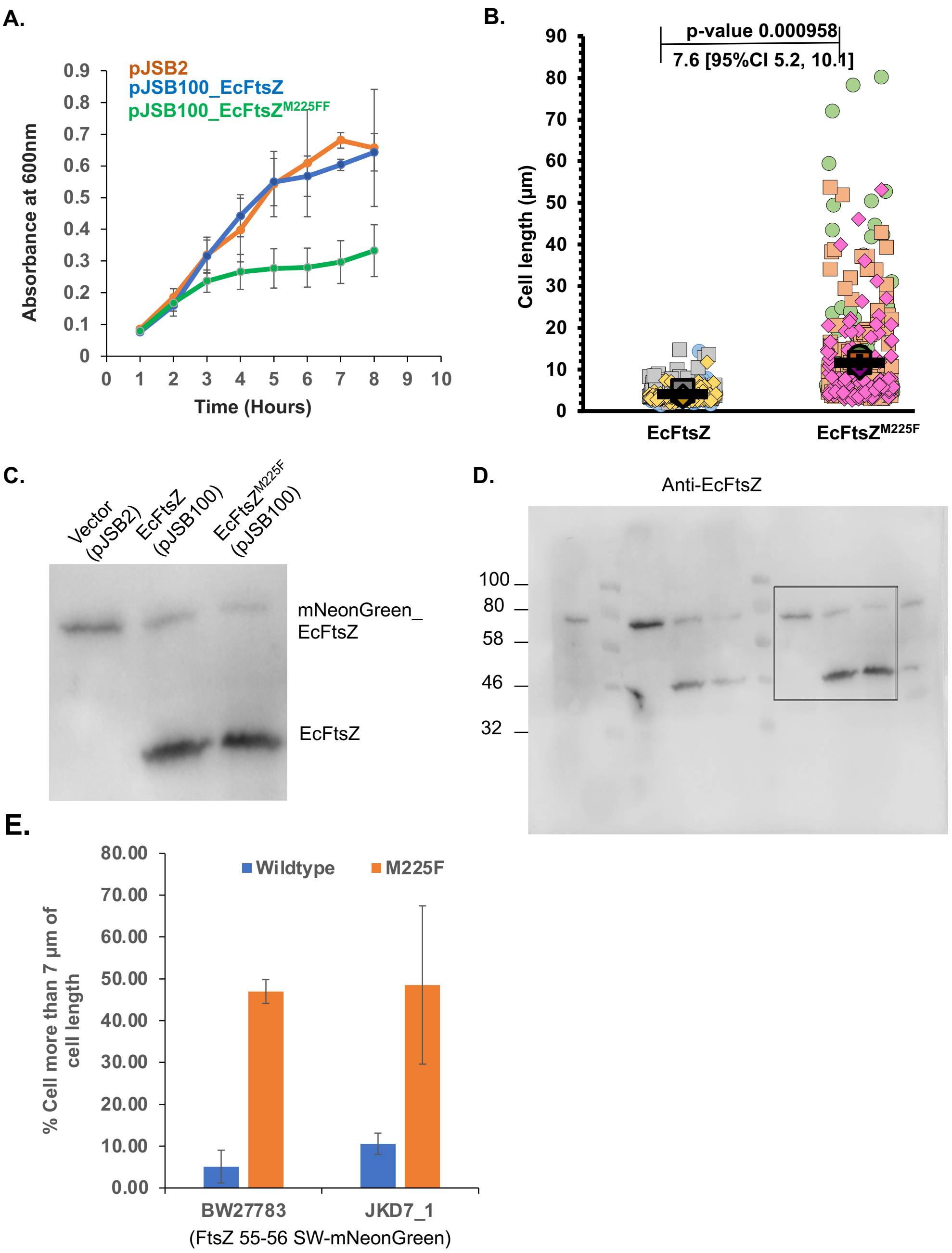


**Figure S4. *In vivo* study comparing wild type and mutant of EcFtsZ.**

**A)** The growth curve showing the growth pattern of the EcFtsZ wildtype and M225F taken in small interval of 1 hr while incubating at 37 °C for 8 hrs. **B)** Cultures of CCD288 strain (BW27783_FtsZ 55-56 mNeonGreen-SW) were grown at 37°C in presence of 0.2% arabinose until they reached an $OD_{600}$ of 0.4-0.6 and imaged by phase-contrast microscopy. The quantitative data of the cell length for EcFtsZ and EcFtsZ$^{M225F}$ expressed in CCD288 are shown as super plots. Expression of the mutant EcFtsZ$^{M225F}$ resulted in an average cell length of 11.73 µm ± 1.56 (number of cells counted were at least 226, N=3), as compared to those expressing wildtype EcFtsZ, which had an average length of 4.09 µm ± 0.81 (number of cells counted were at least 166, N=3). P-values and effect size are indicated above the plots. **C)**, **D)** The part of complete western blot which is showing the expression of EcFtsZ protein expressed through a CCD288 strain having mNeonGreen tagged endogenous FtsZ (band at 63 kDa) and pJSB100 plasmid expressing wildtype and M225F respectively (band at 40 kDa). The first lane represents the strain with empty plasmid expressing just the genomic mNeonGreen tagged EcFtsZ. **E)** Compiled cell length data for both wildtype and M255F mutant of EcFtsZ of CCD288 and JKD7_1 strain representing percentage of cells with cell length more than 7 µm.

Figure S5

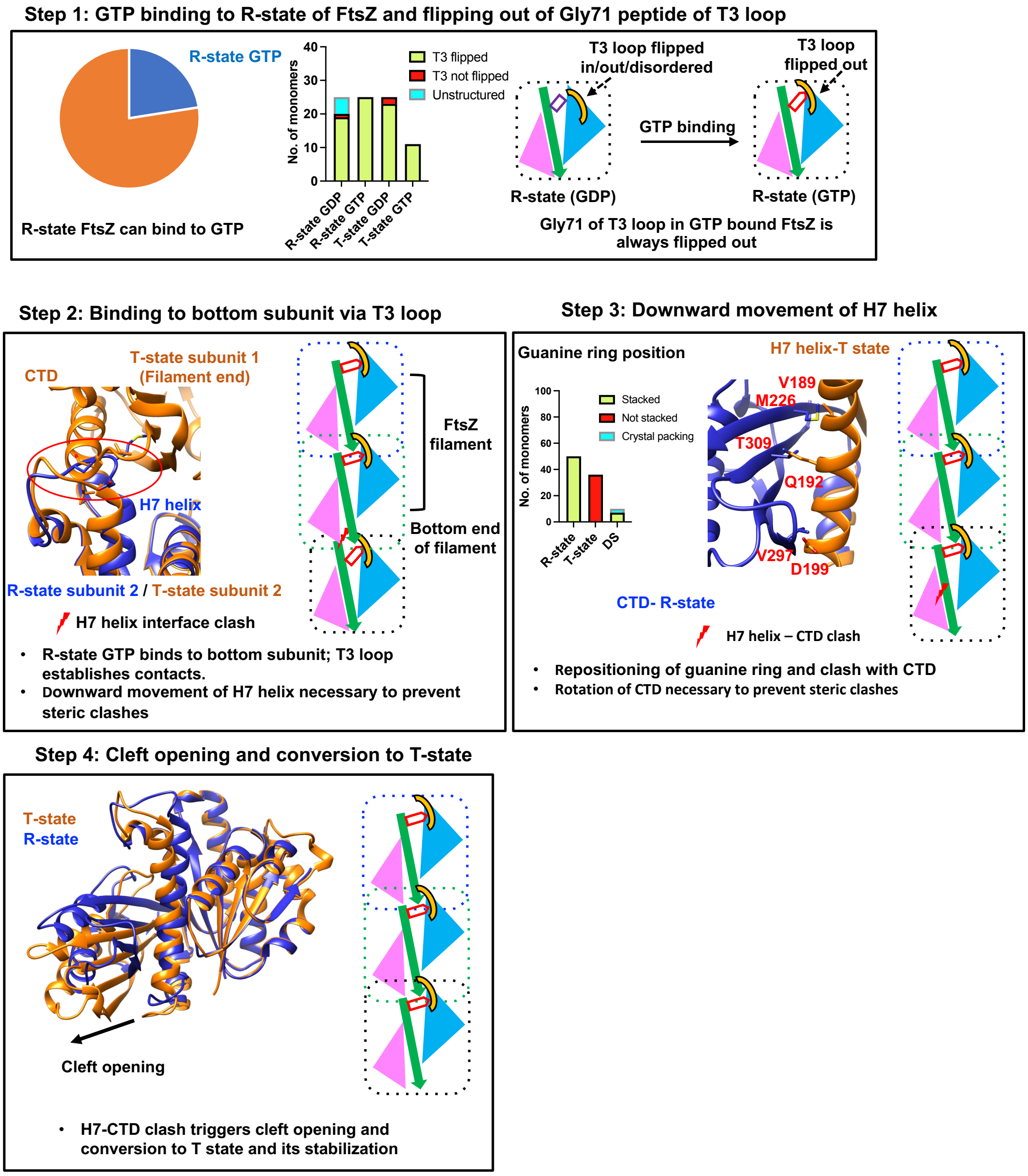


**Figure S5. Model for the kinetic polarity of FtsZ during polymerization**

**Step 1 - GTP binding to monomer FtsZ and flipping out of T3 loop**: The R state of FtsZ can bind to GTP. There are 25 monomers available in PDB, which are captured in the R-state bound to GTP. The GTP binding leads to flipping of the Gly71 peptide and also the gamma phosphate directly interacts with the T3 loop. There are structures of FtsZ with GDP, where the peptide may or may not be flipped out, but all structures of FtsZ with GTP or its analog have flipped out Gly71 peptide.

**Step 2 - Binding to bottom subunit via T3 loop**: The GTP bound R-state FtsZ can, in principle, attach to the bottom interface of the FtsZ filament via the flipped out T3 loop (**Figure 4F**). The H7 helix has to move downwards to clear the steric clash with the top subunit.

**Step 3** - **Downward movement of H7 helix**: H7 helix has to move downwards to clear the steric clash with the top subunit. The downward movement of the H7 helix repositions Phe183, which forms stacking interaction with the guanine ring of the nucleotide. This leads to the repositioning of the guanine ring, which is observed in T-state structures. As the H7 helix moves downwards, there are clashes between the residues of the H7 helix and the beta sheets of CTD.

**Step 4** - **IDC opening and conversion to T state**: The clashes in the interdomain cleft trigger the opening of the cleft and changing the entire molecule to T-state. Note that the H7 helix of the T-state (brown) clashes with the CTD in R-state (deep blue).

The conversion of T-state occurs upon binding to the filament, only after GTP binding. The NTD is in sky blue; the H7 helix and the T7 loop is denoted in green; the CTD is in pink; the purple rectangle denotes GDP and the red pentagon denotes GTP.

### Supplementary tables

#### Table S1: Kinetic parameters:

| Parameters | EcFtsZ | SmFtsZ | SmFtsZ$^{F224M}$ |
|---|---|---|---|
| $E_t$ (μM) | 5 | 10 | 5 |
| $V_{max}$ (μM/min) | 7.64 ± 0.28 | 8.72 ± 0.43 | 8.808 ± 0.34 |
| $k_{cat}$ (min$^{-1}$) | 1.506 ± 0.05 | 0.87 ± 0.04 | 1.76 ± 0.068 |
| $K_M$ (μM) | 35 | 232 | 108 |
| Catalytic efficiency ($k_{cat}$ / $K_M$) (min$^{-1}$ μM$^{-1}$) | 43*10$^{-3}$ | 3.7*10$^{-3}$ | 16*10$^{-3}$ |
| Approximate critical concentration (μM) | 0.5 | 4.1 | 0.8 |

#### Table S2: $k_{obs}$ for all the constructs:

| Constructs | $k_{obs}$ (min$^{-1}$) |
|---|---|
| EcFtsZ | 1.79 ± 0.04 |
| SmFtsZ | 0.62 ± 0.02 |
| SmFtsZ$^{CT}$ | 0.55 ± 0.01 |
| SmFtsZ$^{F224M,\Delta CT}$ | 1.74 ± 0.07 |
| SmFtsZ$^{F224M}$ | 1.52 ± 0.02 |
| SmFtsZ$^{D211G}$ | 0.04 ± 0.01 |

**Table S3: Data collection and refinement statistics:**

| | **SmFtsZ** | **SmFtsZ-GMPPNP** | **SmFtsZ$^{F224M,\Delta CT}$-GDP** |
|---|---|---|---|
| **PDB ID** | **7YSZ** | **7YOP** | **8GRW** |
| **Data Collection statistics** | | | |
| Collected at | DLS, UK | DLS, UK | XRD2, ELETTRA, Trieste |
| Wavelength (Å) | 0.976 | 0.976 | 1 |
| Space Group | $P4_12_12$ | $P4_12_12$ | $P4_12_12$ |
| a, b, c (Å) | 106.59,106.59,128.61 | 106.55,106.55,127.60 | 106.68,106.68,128.78 |
| α, β, γ (˚) | 90, 90, 90 | 90, 90, 90 | 90, 90, 90 |
| Resolution (Å)* | 49.16-2.3 (2.38-2.3) | 49.24-2.2 (2.27-2.2) | 106.67-2.4 (2.49-2.4) |
| Number of unique reflections* | 33121 (3217) | 219453 (17906) | 29504 (3038) |
| $R_{merge}$ (%) * | 8.4 (69.1) | 9 (64) | 9.6 (68.4) |
| $R_{pim}$ (%) * | 4.3 (35.6) | 2.5 (18.3) | 5.7 (41.2) |
| $CC_{half}$* | 0.998 (0.808) | 0.998 (0.924) | 0.998 (0.785) |
| Mean I/σI | 14.9 (3.1) | 21 (4.6) | 15.1 (2.8) |
| Completeness (%) * | 99.7 (100) | 99.9 (98.3) | 99.5 (99.3) |
| Redundancy* | 8.7 (8.8) | 7.5 (7.1) | 7.0 (7.0) |
| **Refinement statistics** | | | |
| Resolution (Å) | 49.16-2.3 | 49.24-2.2 | 49.28- 2.4 |
| Number of unique reflections (test set) | 33090 (1657) | 38229 (1915) | 29477 (1503) |
| $R_{work}$ / $R_{free}$ (%) | 0.179/0.235 | 0.173/0.221 | 0.192/0.236 |
| Average B-factor (Å$^2$) | 50 | 43 | 46 |
| Wilson B-factor (Å$^2$) | 41.8 | 33.4 | 38.7 |

| | | | |
|---|---|---|---|
| RMS deviations | | | |
| Bond lengths (Å) | 0.006 | 0.011 | 0.01 |
| Bond angles (˚) | 0.827 | 1.148 | 0.972 |
| Ramachandran map statistics | | | |
| Favoured (%) | 96.9 | 98.30 | 97.92 |
| Allowed (%) | 3.1 | 1.70 | 2.08 |
| Outliers (%) | 0 | 0 | 0 |

*Values in parentheses denote last resolution shell

**Table S4: FtsZ structures for analysis**

| T state | T state | R state | R state | DS |
|---|---|---|---|---|
| 3WGJ | 7OMQ | 5MN5 | 6UNX | 7YOP |
| 4DXD | 7OJZ | 3WGL | 6UMK | 7YSZ |
| 5H5H | 7OI2 | 3WGK | 6LL6 | 8GRW |
| 3VO8 | 7ON2 | 5H5I | 6LL5 | 1W5F |
| 3VOA | 7ON3 | 5MN6 | 5V68 | 5MN7 |
| 5MN4 | 7ON4 | 5MN8 | 5ZUE | |
| 3VOB | 7OJA | 4U39 | 1W5A | |
| 3WGN | 7OJB | 2VXY | 1W5B | |
| 5XDT | 7OJC | 2RHL | 1W5E | |
| 5XDU | 7OJD | 2RHJ | 1W58 | |
| 5XDW | 5XDV | 2RHO | 1FSZ | |
| 3WGM | 6KVP | 2VAM | 2VAP | |
| 5H5G | 6KVQ | 2RHH | 1W59 | |
| 7OHH | 6SI9 | 2Q1X | 4E6E | |
| 7OHK | | 4KWE | 2R6R | |
| 7OHL | | 1RQ2 | 2R75 | |
| 7OHN | | 1RQ7 | 1OFU | |
| 7OMJ | | 1RLU | 2VAW | |
| 7OMP | | 2Q1Y | | |

**Table S5: % of cells above 7 um cell length of wild type and cleft mutant**

| Strain | EcFtsZ | $EcFtsZ^{M224F}$ |
|---|---|---|
| BW27783 | 5.10 ± 3.92 | 46.98 ± 2.84 |
| JKD7_1 | 10.56 ± 2.55 | 48.54 ± 18.93 |

**Table S6: List of strains**

| Strains | |
|---|---|
| CCD57 | JKD 7-1/ pKD3 (ftsZ)/ pJSB2 |
| CCD58 | JKD7-1/ pKD3/ pJSB100_EcFtsZ |
| CCD288 | BW27783 FtsZ 55-56 SW-mNG |
| CCD161 | BW27783 |
| CCDE603 | JKD7_1/pKD3/pJSB100_EcFtsZ_M225F_untagged |
| CCDE497 | BW27783 FtsZ 55-56 SW-mNeonGreen/pJSB2 |
| CCD359 | BW27783 FtsZ 55-56 SW-mNeonGreen/pJSB100_EcFtsZ |
| CCDE599 | BW27783 FtsZ 55-56 SW-mNeonGreen/pJSB100_EcFtsZ_M225F |
| CCD252 | JS964; *ΔminCDE* |
| CCDE668 | JS964(delMinCDE)/ pJSB100_EcFtsZ_mNeongreen_MinD_mts |
| CCDE669 | JS964(delMinCDE)/ pJSB100_SmFtsZ_core_mNeongreen_MinD_mts |
| CCDE667 | BW27783/pJSB100_EcFtsZ_mNeongreen_MinD_mts |

**Table S7: List of plasmids**

| Plasmid number | name |
|---|---|
| pCCD434 | TG1/ pJSB2 |
| pCCD436 | TG1/ pJSB100EcFtsZ |
| pCCD907 | DH10β/pJSB100-EcFtsZ1-366-Linker-mNeonGreen-EcMinD_MTS256–270 |
| pCCD1020 | DH10β/pJSB100_EcFtsZ_M225F_untagged |
| pCCD1042 | DH10β/pJSB100_SmFtsZ_full_length |
| pCCD1043 | DH10β/pJSB100_SmFtsZ_core_314a.a_mNeongreen_MinD_mts |
| pCCD1044 | DH10β/pJSB100_SmFtsZ_F224M_core_314a.a_mNeongreen_MinD_mts |

**Table S8: List of primers**

| Primer name | Sequence (5' to 3') | Purpose |
|---|---|---|
| EcFtsZ_F | ATGTTTGAACCAATGGAACTTAC | EcFtsZ gene-specific amplification |
| EcFtsZ_R | ATCAGCTTGCTTACGCAGGAATG | EcFtsZ gene-specific amplification |
| EcFtsZ_vF | GTTTAACTTTAAGAAGGAGATATAC ATATGTTTGAACCAAT GGAACTTACC | EcFtsZ clone; primer with vector overhang for incorporation into plasmid |
| EcFtsZ_vR | GCTTTTAGTGGTGATGGTGATGAT GGG ATCCATCAGCTTGCTTACGC AGGAATGC | EcFtsZ clone; primer with vector overhang for incorporation into plasmid |
| SmFtsZ_F | ATGGACAATTTTGATAATTATGAAC AAGTCGCG | SmFtsZ gene-specific amplification |
| SmFtsZ_R | CAACTACGACGAACAAATGGTGGT AA ATCATC | SmFtsZ gene-specific amplification |
| SmFtsZ_vF | GTTTAACTTTAAGAAGGAGATATAC ATAT GGACAATTTTGATAATTATG | SmFtsZ clone; primer with vector overhang for incorporation into plasmid |
| SmFtsZ_vR | CTTTTAATGATGATGATGATGATGG GAT CCCCAACTACGACGAACAAATG | SmFtsZ clone; primer with vector overhang for incorporation into plasmid |
| SmFtsZ_384 F | CGAATTAATGCGTGGCGAGAGCAT G | SmFtsZ clone; To replace TGA codon |

| | | |
|---|---|---|
| | TTAATAATAATC | |
| SmFtsZ$^{\Delta CT}$ _R | GCTTTTAATGATGATGATGATGATG G GATCCATCAAACCCAGTCGC | To generate SmFtsZ$^{\Delta CT}$ |
| SmFtsZ$^{D211G}$_ F | CCTTAATTAATTTAGACTTTGCAGG TA TTAAGACTGTTATG | To generate SmFtsZ$^{D211G}$ in SmFtsZ$^{\Delta CT}$ background |
| SmFtsZ$^{F224M}$ _F | GGGAACGCTTTAATGGGGATCGG TATTGG | To generate SmFtsZ$^{F224M}$ and SmFtsZ$^{F224M\Delta CT}$ |
| RSO 1081 pJSB100_S mFtsZ_F | CGTTTTTTTGGGCTAGCGGAGTGC ACCCTATGGACAATTTTGATAATT ATGAAC | To generate SmFtsZ in pJSB100 and pJSB100_mNeonGreen_ MinD_mts plasmids |
| RSO 1083 pJSB100_S mFtsZ_R | CATGCCTGCAGGTCGAGGTACCC GATCCTTATTACCAACTACGACGA ACA | To generate SmFtsZ in pJSB100 plasmid |
| RSO 1082 MinD_mts_ mNeonGree n_linker_Sm FtsZ_R | CTGACGCGGCCGCCTGCTGGCCT AGGTGGCCCAGTCGCAATTACAG TTA | To generate SmFtsZ in pJSB100_mNeonGreen_ MinD_mts plasmid |
| RSO 1107 EcFtsZ_M22 5F_F | CTGAGATGGGCTACGCAATGATG GGTTCTGGCGTGGCGAGCGGTG | To generate SmFtsZ_M225F in pJSB100 plasmid |
| RSO 428 pBAD33_R | GATTTAATCTGTATCAGG | To generate SmFtsZ_M225F in pJSB100 plasmid |

**Table S9: Crystallization conditions**

| | **PDB ID: 7YSZ** | **PDB ID: 7YOP** | **PDB ID: 8GRW** |
|---|---|---|---|
| Method | Sitting drop vapour diffusion | Sitting drop vapour diffusion | Hanging drop vapour diffusion |
| Plate type | 48 | 48 | 24 |
| Temperature (K) | 291 | 291 | 291 |
| Construct and protein concentration (mg/ml) | SmFtsZ$^{\Delta CT}$ - 10mg/ml | SmFtsZ$^{\Delta CT}$ - 10mg/ml | SmFtsZ$^{F224M\Delta CT}$ - 10mg/ml |
| Buffer composition of protein solution | 50mM HEPES pH= 6.5, 50mM KCl | 50mM HEPES pH= 6.5, 50mM KCl, 2mM GMPPNP, 5mM MgCl2 | 50mM HEPES pH= 6.5, 50mM KCl |
| Composition of reservoir solution | 1.5M ammonium sulphate, 0.1M Tris pH= 8.5 | 0.3M sodium acetate, 0.1M Tris pH= 8.5, 15% w/v PEG 4000 | 0.3M sodium acetate, 0.1M Tris pH= 8.5, 15% w/v PEG 4000 |
| Volume and ratio of drop | 500 nl : 500 nl | 500 nl : 500 nl | 2.5 µl : 2.5 µl |
| Volume of reservoir (ml) | 0.1 | 0.1 | 0.75 |
